# Closed-loop stimulation of the medial septum terminates epileptic seizures

**DOI:** 10.1101/2020.03.09.982827

**Authors:** Yuichi Takeuchi, Márk Harangozó, Lizeth Pedraza, Tamás Földi, Gábor Kozák, Antal Berényi

## Abstract

Temporal lobe epilepsy with distributed hippocampal seizure foci is often intractable and its secondary generalization might lead to sudden death. Early termination through spatially extensive hippocampal intervention is not feasible directly, due to its large size and irregular shape. In contrast, the medial septum (MS) is a promising target to govern hippocampal oscillations through its divergent connections to both hippocampi. Combining this ‘proxy intervention’ concept and precisely timed stimulation, we report here that closed-loop MS electrical stimulation can quickly terminate intrahippocampal seizures and suppress secondary generalization in a rat kindling model. Precise stimulus timing governed by internal seizure rhythms was essential. Cell-type-specific stimulation revealed that precisely timed activation of MS GABAergic neurons underlay the effects. Our concept of phase-targeted proxy stimulation for intervening pathological oscillations can be extrapolated to other neurological and psychiatric disorders, and its current embodiment can be directly translated into clinical application.

---

Approximately 1% of world population have epilepsy and 30% of them are still refractory (1, 2). Temporal lobe epilepsy (TLE) is often drug-resistant and uncontrolled TLE can lead to secondary generalized seizures and consequently to sudden death (3, 4). Although effective, surgical resection of the seizure focus is highly invasive, irreversible, and frequently associated with cognitive dysfunctions and it may not be the best solution for patients with ambiguous or multifocus bilateral TLE (5–7). Thus, a new therapeutic approach is desired. Deep brain stimulation (DBS) has been employed to treat neurological and psychiatric disorders including epilepsy (8). Direct stimulation of the hippocampus (HPC) can reduce seizure occurrence especially in medial TLE (8, 9). However, because the HPC is a relatively large bilateral structure, direct electrical stimulation may affect only a small portion of the seizure generating structure (10). An alternative therapeutic approach is stimulation of another brain region than the seizure focus (11–13).

Here, we propose that activity of the HPCs can be thoroughly controlled in a technically feasible way by interfering with a smaller circumscribed structure, which has massive divergent efferent connections to the whole extent of HPCs (proxy intervention). The medial septum (MS) is an attractive target of such proxy intervention because it is a small midline structure with diverging projections onto both HPCs (14). The septohippocampal pathway plays an important role in governing theta (15, 16) and high-frequency HPC oscillations (17); the latter are particularly vulnerable to become seizure activity (18–20).

Clinical DBS applications, either being free-running or responsive (i.e. triggered by the detection of certain activity patterns), deliver hard-coded, non-adaptive sequences of pre-set pulse trains or random noise waveforms (21). Although these open-loop approaches are effective in reducing seizure occurrence by decreasing seizure susceptibility, they cannot quickly terminate already initiated seizures (22). This would require temporally aligned stimulus delivery targeting specific phases of pathological oscillations (23–25). Due to its direct reciprocal connections, the septohippocampal axis is resonating in a highly coherent manner in physiological and pathophysiological conditions including ictal periods of epilepsy (26, 27). Stimulation of MS neurons can precisely entrain HPC local field potentials (LFPs) (28–30). In addition, the MS can powerfully gate oscillations not only in the HPC but also in the entorhinal cortex (EC), which is the bottle-neck structure for seizure propagation from the HPC to the neocortex (Ctx) (31). Given these anatomical and temporal features of the septohippocampal axis, we hypothesized that the MS can be a powerful seizure regulation center for on-demand control of TLE. Thus, we implemented a closed-loop stimulation of the MS driven by seizure patterns, and investigated whether phase-targeted precise stimulus delivery to the MS can cease already initiated HPC seizures, before they generalize.

The present study is also designed to obtain a mechanistic understanding of possible seizure terminating effects of the MS in a cell-type specific manner. The MS has GABAergic, glutamatergic (Glut), and cholinergic neuron populations, and all of them project to distinct targets of the HPC and the EC (32–34), with distinct excitatory or inhibitory postsynaptic effects on these targets (29, 34, 35). Optogenetics has been successfully applied to the MS populations to dissect their physiological roles in the septohippocampal axis (17, 29, 30). We employed selective optogenetic effectors in our closed-loop approach to dissect the hypothesized seizure terminating effects.

Here, we found that seizure rhythm congruent MS electrical stimulation can effectively terminate HPC seizures in their early phase and suppress secondary generalization. Responsive, but open-loop (fixed pattern) MS electrical stimulation did not terminate the seizures, which emphasized that precise seizure phase-matched stimulus delivery is essential to break ongoing pathologic oscillations. Time-targeted optogenetic experiments revealed that precisely timed activation of MS GABAergic neurons mainly underlay the seizure termination effect, while pre-ictal activation of MS cholinergic neurons can reduce seizure susceptibility.

## Results

### Responsive MS stimulation for intervening seizures of HPC-origin

HPC kindled rats were employed as a model of refractory TLE (36); rats were fully kindled by daily electrical stimulation (E-stim) of the HPC commissure (n = 25). In the kindled rats, the commissure stimulation instantaneously induced massive 14–20 Hz after-discharges in the bilateral HPCs (Figure 1A, B). The HPC seizures were then secondary generalized, resulting in Racine’s scale 4 or 5 motor seizures (Figure 1A, C; Dataset figure 1, see Table S2). To terminate these, we focused on two types of responsive MS E-stim: open-loop and closed-loop interventions (Figure 1D, E). For open-loop intervention, stimulus trains at fixed rates were delivered in response to seizure episodes. For closed-loop intervention, each stimulus pulse was triggered by seizure waves themselves, meaning that the MS was stimulated with the internal, real-time rhythmicity of HPC seizures (closed-loop seizure rhythm stimulation). A delay of 0 to 60 ms was introduced to shift the stimulus pulses to target specific phases of the seizure waves (23).

**Figure 1.**
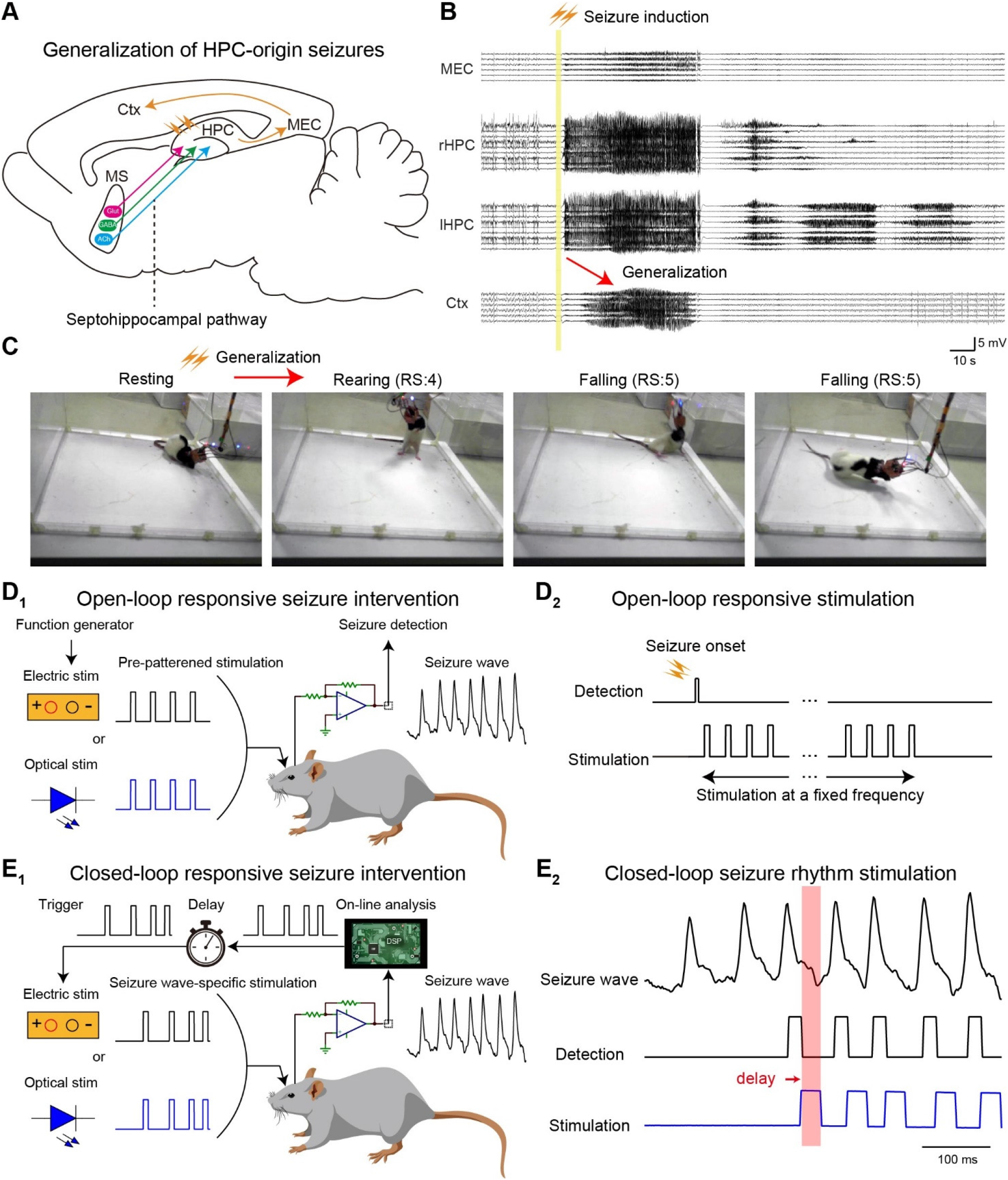
Responsive MS stimulation for intervening seizures of HPC-origin. (**A**) Schema of generalization of HPC seizures and septohippocampal pathway. MEC, medial entorhinal cortex. (**B**) Representative LFP traces of HPC seizures propagated to the Ctx. (**C**) E-stim of the HPC commissure induced motor seizures as well. RS: the Racine’s scale. (**D**_**1**_, **D**_**2**_) The concept of open-loop responsive intervention. (**E**_**1**_) The concept of closed-loop responsive intervention (seizure rhythm-driven stimulation). Seizure waves were detected in real-time and each detection triggers MS stimulation with a defined delay. (**E**_**2**_) A representative HPC seizure, its detection, and MS stimulation. Red stripe indicates the targeted phase. See also Figure S1.

### Open-loop MS E-stim has no effects or promotes HPC seizures

Open-loop responsive MS E-stim was examined first (Figure 2A). Each MS E-stim (±400 μA biphasic, 1 ms-long) evoked robust and transient potentials in the HPC and the EC (1–4 mV) but not in the Ctx (Figure S1A, C). Following the seizure induction, MS E-stim trains were delivered at 1, 8, or 20 Hz for two minutes (Figure 2B); those stimuli themselves did not cause any seizures in non-kindled rats (data not shown). The open-loop MS E-stim did not shorten HPC electrographic seizures at any frequencies examined (50.5 ± 19.9 s to 53.9 ± 26.2 s, P > 0.05, paired t-test, Figure 2C, E; Trial numbers, animal numbers, and other statistics are fully reported in the figure legends and in the Table S1). Modulation indices (MIs) of the HPC seizures by MS E-stim were distributed around 0 and its distribution was not skewed, which meant almost no modulation (*P* = 0.39, two-sample Kolmogorov-Smirnov test, Figure 2D) Neither Ctx electrographic nor motor seizures were alleviated by the stimulation (28.3 ± 19.4 s to 25.2 ± 20.0 s and 4.2 ± 1.4 to 4.1 ± 1.6 in Racine’s scale, *P* = 0.37 and 0.47, paired *t*-test and Wilcoxon singed rank test, respectively; Figure 2C, E).

**Figure 2.**
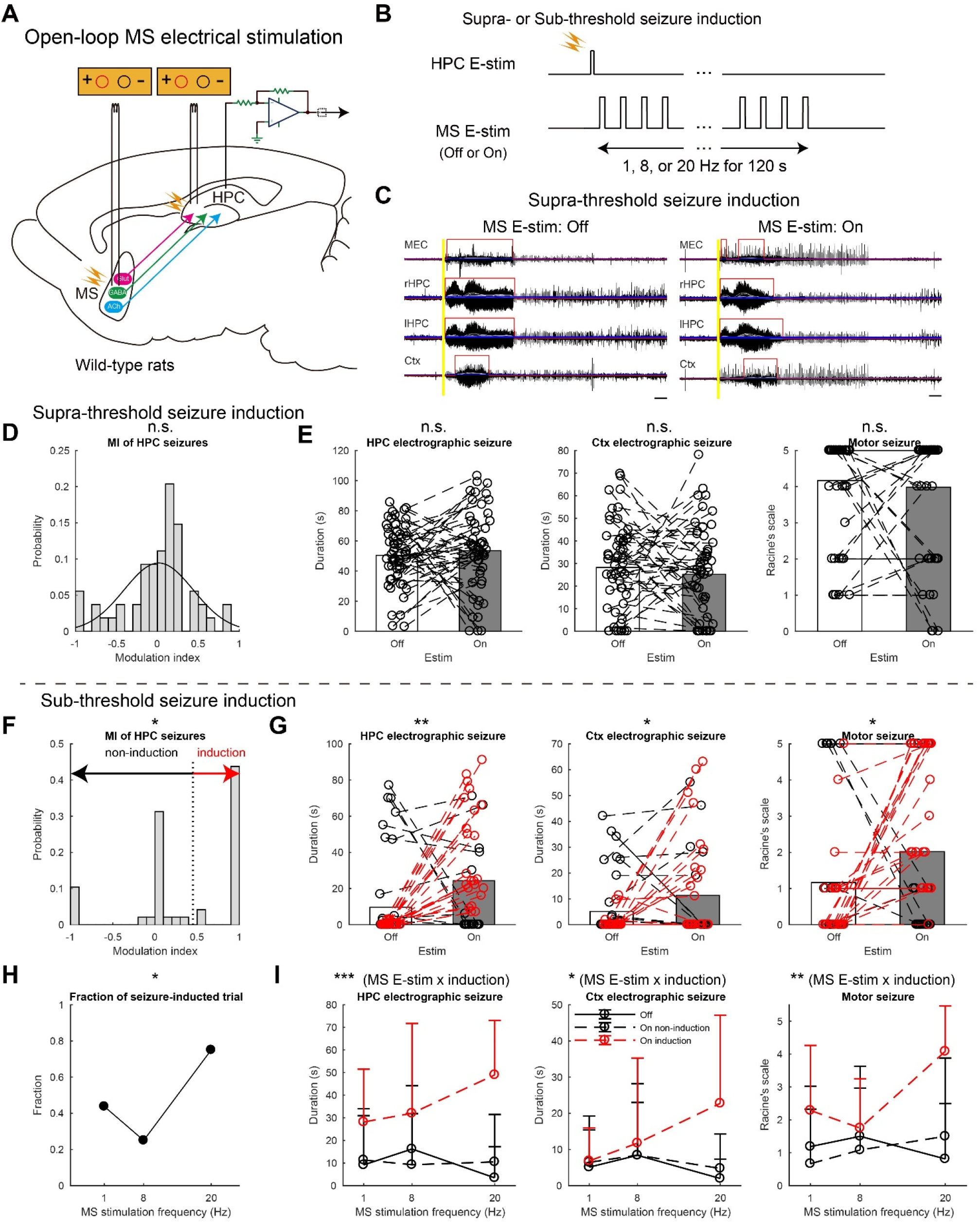
Open-loop MS E-stim has no effects or promotes HPC seizures. (**A**) Schema of the experiment. Stimulating electrodes were inserted into the HPC and the MS for seizure induction and intervention, respectively. (**B**) Open-loop responsive intervention. Seizures were induced at supra- or sub-threshold intensity. (**C**) Representative seizure waves induced by the supra-threshold stimulation w/wo open-loop MS E-stim. Six to nine 10–80 Hz bandpass-filtered LFP traces were overlapped for each brain region. Grey traces represent smoothed LFP traces with a moving average filter. Each blue line represents three rms level of baseline of each smoothed LFP trace. Red lines represent detected seizures. Scale bars: 10 s. (**D**) Distribution of modulation indices (MIs) of duration of HPC seizures by MS E-stim. A black curve represents a fitted Gaussian model. (**E**) Duration of HPC and Ctx seizures, and motor seizures. Data of different MS stimulus frequencies were pooled. Each marker pair with a dashed line represents a pair of consecutive non-stimulated and stimulated trials with the same condition. Values are represented as bars for means and circles for each trial. (**F**) Distribution of MIs. Each trial pair was clustered into non-seizure induction or seizure induction group by the absolute value of the global threshold was determined via the whole data set (see Methods, and Dataset figure 2). (**G**) The same conventions as (E) but with the clustered data. Red colored data represent the seizure-induction cluster. (**H**) Fractions of seizure-induced trials against MS E-stim frequencies. (**I**) MS E-stim frequency resolved representation of the data shown in (G). Values are represented as means ± s.d. n = 108 and 96 trials from four rats for (D, E) and (F–I), respectively. Statistical significance was tested by two-sample Kolmogorov-Smirnov for asymmetry (skewness) of MI distribution (D, F), paired *t*-test for seizure durations and Wilcoxon singed rank test for the Racine’s scale (E, G), χ^2^ test for (H), and three-way repeated ANOVA for (I). Results of statistical tests are extensively reported in Table S1. n.s., not significant; \**P* < 0.05; \*\**P* < 0.01, \*\*\**P* < 0.001. See also Figure S2.

We then examined effects of this open-loop MS E-stim on incomplete seizure episodes evoked by sub-threshold commissure stimulation, which typically resulted in non-generalized partial seizures (Figure 2F–I). The open-loop MS E-stim significantly elongated HPC seizures (9.6 ± 22.1 s to 24.3 ± 28.5 s, *P* < 0.01, paired *t*-test, Figure 2G). To underpin a factor for this pro-seizure effect, stimulated and non-stimulated trial pairs were classified into seizure-induction (MI > 0.5) and non-seizure-induction trials (MI ≤ 0.5) (*P* < 0.05, two-sample Kolmogorov-Smirnov test, Figure 2F). The fraction of seizure-induction trials was significantly higher at 20 Hz (0.750) than those at 1 Hz and 8 Hz (0.438 and 0.250, respectively; *P* < 0.05, χ^2^ test, Figure 2H). The open-loop MS E-stim significantly deteriorated Ctx and motor seizures (5.6 ± 10.9 s to 11.8 ± 18.9 s and 1.2 ± 1.9 to 2.0 ± 2.1 in Racine’s scale, *P* < 0.05 for both, paired *t*-test and Wilcoxon signed rank test, respectively; Figure 2G). Notably, 20 Hz stimulation induced longer Ctx seizures (22.8 ± 24.4 s) and higher Racine’s scale values (4.1 ± 1.4) in seizure-induction trials (Figure 2I). It is worthy to note that open-loop MS E-stim at higher frequency (≥ 40 Hz) induced generalized episodes with Racine’s scale 4 or 5 motor seizures even in non-kindled rats (data not shown). These results suggest that open-loop MS E-stim does not alleviate HPC seizures and a false positive triggering of an open-loop responsive stimulator can even make seizures worse.

### Closed-loop seizure rhythm-driven MS E-stim terminates seizures of HPC-origin

MS E-stim was then delivered in a closed-loop manner (Figure 3A, B). The induced HPC seizures were processed with a custom-made algorithm in real-time and each HPC LFP deflection of the 10–130 Hz range triggered a single-pulse MS E-stim (±400 μA biphasic, 1 ms-long) with a fixed delay. In contrast to the open-loop stimulation, closed-loop seizure rhythm-driven MS E-stim significantly shortened HPC seizures (53.6 ± 22.3 s to 31.8 ± 25.8 s, *P* < 0.001, paired *t*-test, Figure 3C, D, F). The closed-loop stimulation also shortened Ctx seizures (28.1 ± 1.51 s to 13.2 ± 17.0 s, *P* < 0.001, paired *t*-test) and decreased seizure severity according to Racine’s scale (4.4 ± 1.1 to 2.1 ± 2.3 in Racine’s scale, *P* < 0.001, Wilcoxon signed rank test). Because these seizure terminating effects were rather in an all-or-none manner, distribution of MIs of HPC seizures could be fitted with a two-component Gaussian mixture model and off-on trial pairs were classified into success and non-success trials (*P* < 0.001, two-sample Kolmogorov-Smirnov test, Figure 3E). Fractions of success trials were not significantly different between delays (0.615, 0.500, 0.571, and 0.688 for 0, 20, 40, and 60 ms, respectively; *P* = 0.766, χ^2^ test, Figure 3G). The success and non-success classifications were consistently valid for the Ctx and motor seizures, too; interaction between MS E-stim and success-labelling were all significant on HPC, Ctx, and motor seizures (*P* < 0.001 for all, three-way repeated ANOVA, Figure 3H). On success trials, the average duration of Ctx seizures and Racine’s scale values were robustly reduced (< 4.4 s and < 1.1 in Racine’s scale, Figure 3H), which meant secondary generalization was effectively suppressed. These results indicate strong seizure terminating effects of the closed-loop MS E-stim.

**Figure 3.**
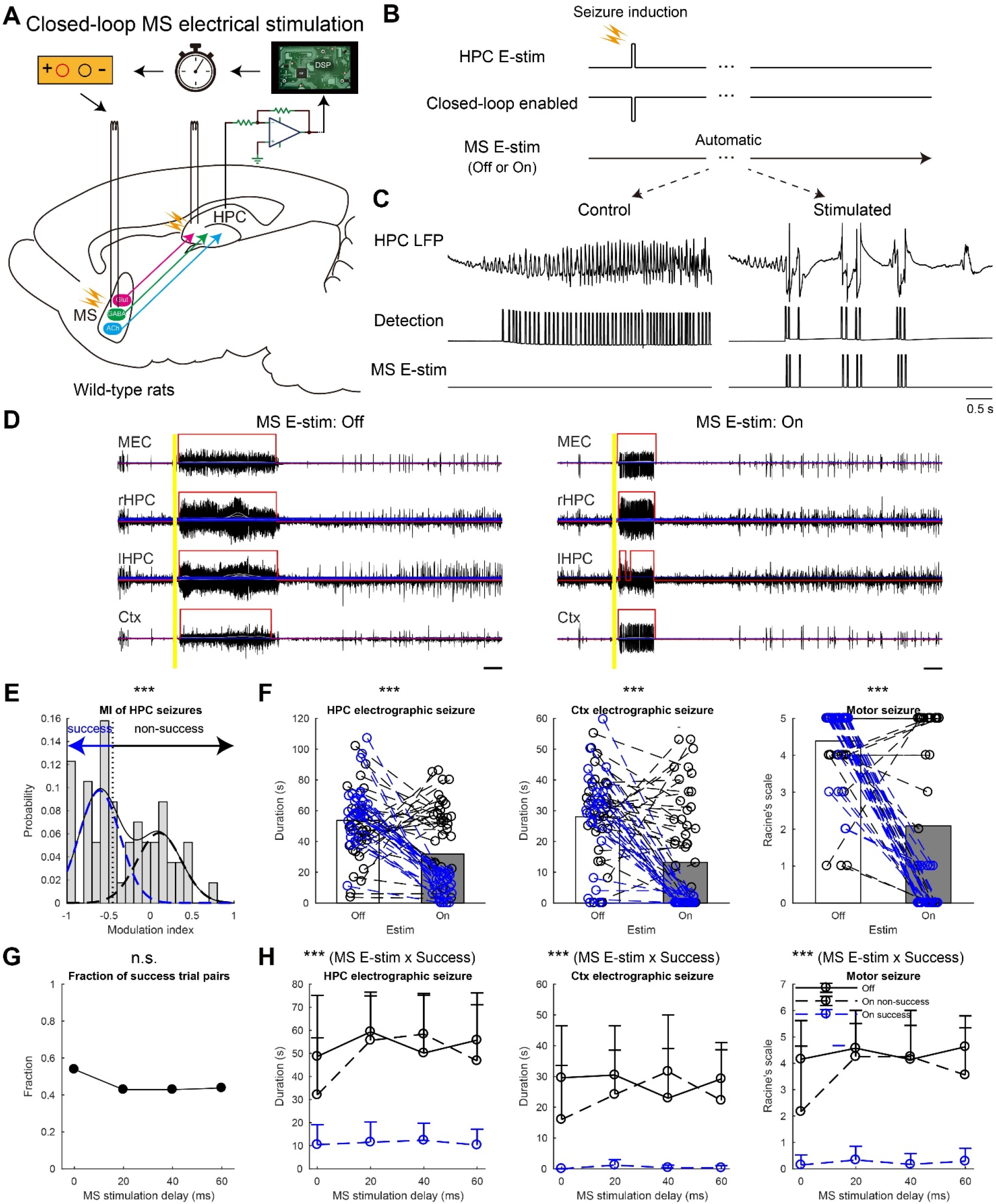
Closed-loop seizure rhythm-driven MS E-stim terminates seizures of HPC-origin. (**A**) Schema of the experiment. (**B**) Close-loop seizure rhythm intervention. Seizures were induced by supra-threshold stimulation. Automatic seizure detection was always turned on except seizure-induction time. Each detection triggered a MS E-stim pulse with a fixed delay. (**C**) HPC seizure waves w/wo the MS E-stim. (**D**) Representative seizure waves w/wo the MS E-stim. The same convention is used as on Figure 2C. (**E**) Distribution of MIs. The distribution was fitted to a Gaussian mixture model and each trial pair was clustered into success or non-success group by the global threshold. (**F**) Duration of HPC and Ctx seizures, and motor seizures with the clusters defined in (E): Blue, success trial pairs; Black, non-success trial pairs. Data with different delays were pooled. (**G**) Fractions of success trial pairs as a function of delays. (**H**) Delay time resolved representation of the data shown in (F). n = 114 trials from three rats. Other conventions for data presentation and statistical tests employed are the same as on Figure 2. n.s., not significant; \*\*\**P* < 0.001. See also Figure S3.

**Figure 4.**
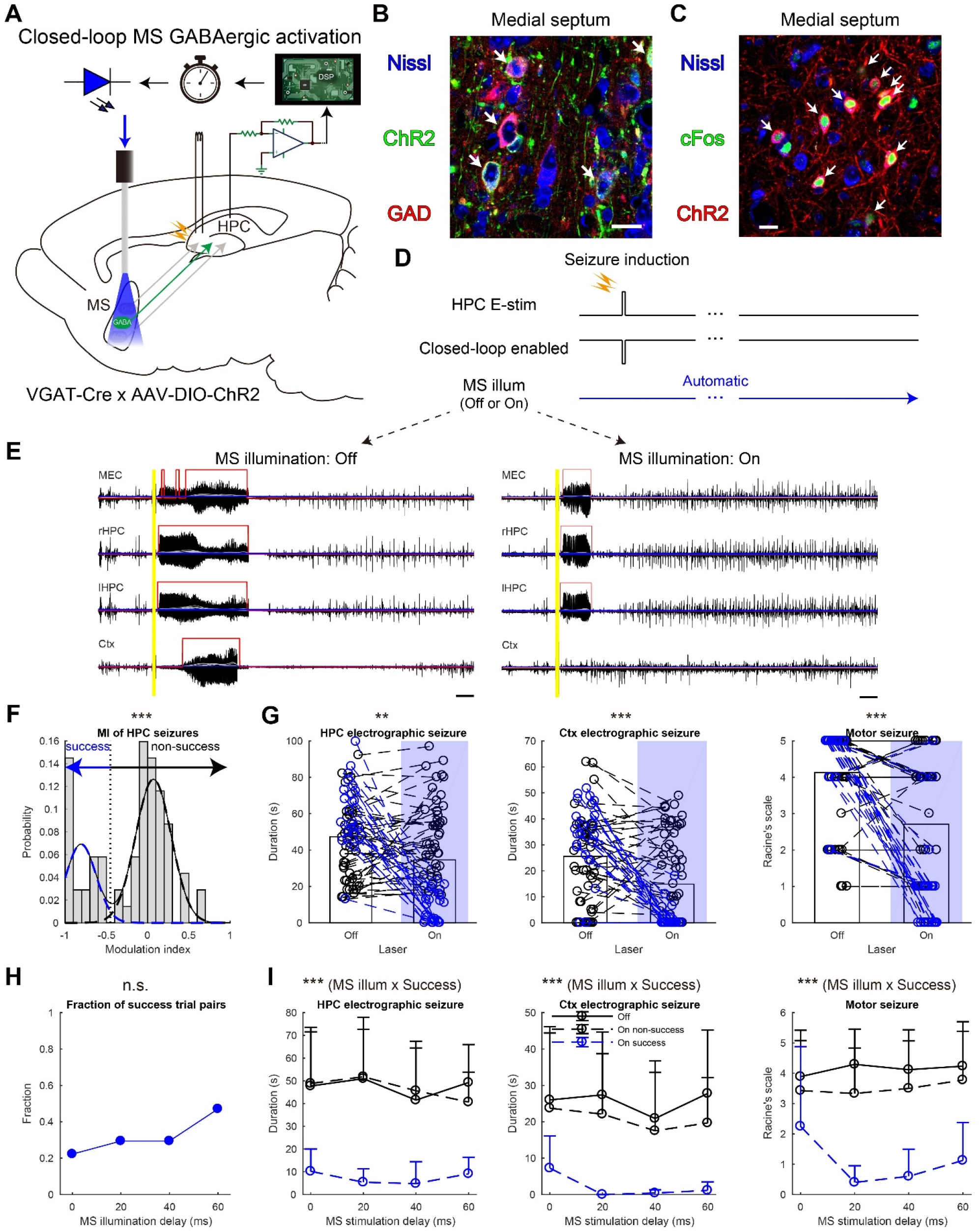
Closed-loop seizure rhythm-driven activation of MS GABAergic neurons terminates seizures of HPC-origin. (**A**) Schema of the experiment. (**B**) ChR2 signals were colocalized with GABAergic neurons. Arrows indicate colocalizations. Scale bar: 20 μm. (**C**) cFos expression was colocalized with ChR2 expressing neurons after illumination. Arrows indicates colocalizations. Scale bar: 20 μm. (**D**) Closed-loop seizure rhythm intervention. Each detection triggered a 30 ms-long MS illumination with a fixed delay. (**E**) Representative seizures w/wo the MS illumination. (**F**) Distribution of MI with the MS illumination. The distribution was fitted with a Gaussian mixture model and each trial pair was clustered into success or non-success trial group by the global threshold. (**G**) Duration of HPC and Ctx seizures, and motor seizures with the clusters defined in (F). (**H**) Fractions of success trial pairs as a function of delays. (**I**) MS illumination delay resolved representation of the data shown in (**G**). n = 138 trials from four rats. Other conventions are the same as on Figure 3. \*\**P* < 0.01; \*\*\**P* < 0.001. See also Figure S4.

### Seizure terminating effects of closed-loop seizure rhythm-driven MS stimulation are mediated by MS neurons

MS E-stim might activate not only MS neurons but also MS targeting HPC neurons via antidromic spikes, nearby brain regions by current spread, and even glial cells (26, 37). To investigate if the seizure terminating effects are mediated specifically by MS neurons, we employed optogenetic stimulation (Dataset figures 3–5). Channelrhodopsin-2 (ChR2) was transduced to MS neurons using a viral vector with a synapsin promoter (Figure S2A). The gene transduction was confined to the MS (Figure S1B), and ChR2 signals were well colocalized with neuronal marker NeuN immunoreactions (Figure S2B). MS blue-light illumination induced cFos expression in ChR2 expressing neurons and modulated unit activities of MS neurons, which meant successful activation of MS neurons (Figure S2C; Dataset figures 5, 6). Each MS illumination reliably induced robust and fast LFP deflections in the HPC and the EC but not in the Ctx (Figure S1D). Similarly to E-stim, open-loop MS illumination did not alleviate HPC seizures (43.6 ± 22.4 s to 40.8 ± 18.9 s, *P* = 0.51, paired *t*-test, Figure S2A, D, E, G, H). Distribution of MI was not skewed (*P* = 0.09, two-sample Kolmogorov-Smirnov test Figure S2F). Neither Ctx nor motor seizures were affected by the open-loop illumination (32.1 ± 14.1 s to 30.9 ± 14.5 s and 4.5 ± 0.9 to 4.3 ± 1.2 in Racine’s scale, *P* = 0.68 and 0.39, paired *t*-test and Wilcoxon signed rank test, respectively; Figure S2G, H). These results further indicate that open-loop MS stimulation is not effective to terminate seizures of HPC-origin.

Closed-loop seizure rhythm-driven activation of non-specific MS neurons was next examined in the same rats (Figure S3A, B). In contrast to the open-loop illumination, closed-loop seizure rhythm MS illumination significantly shortened HPC seizures (48.2 ± 16.6 s to 33.0 ± 25.5 s, *P* < 0.001, paired *t*-test, Figure S3C, E). The illumination also shortened Ctx seizures and decreased Racine’s scale values (34.9 ± 20.0 s to 23.0 ± 20.2 s and 4.2 ± 1.3 to 3.0 ± 2.0 in Racine’s scale, *P* < 0.001 for both, paired *t*-test and Wilcoxon signed rank test, respectively; Figure S3C, E). The MI distribution was well fitted to a two-component Gaussian mixture model; One of the Gaussians was centered around −1 and the other was around zero (*P* < 0.01, two-sample Kolmogorov-Smirnov test, Figure S3D). Illuminated and non-illuminated trial pairs were then classified into success and non-success trial pairs based on the model (Figure S3D). This classification was consistent with all seizure parameters examined (*P* < 0.001, three-way ANOVA, Figure S3G). Manifestations of seizures were significantly alleviated in success trials (< 3.9 s, < 5.4 s, and < 1.9 in Racine’s scale for HPC seizures, Ctx seizures, and motor seizures, respectively; Figure S3G). Fractions of success trials were not significantly different between delays (0.471, 0.444, 0.125, and 0.267 for 0, 20, 40, and 60 ms delay, respectively; *P* = 0.12, χ^2^ test, Figure S3F). Thus, closed-loop seizure rhythm activation of MS neurons has seizure terminating effects compatible to those of MS E-stim. These results suggest that the seizure terminating effects of closed-loop seizure rhythm-driven MS E-stim are presumably mediated by MS neurons.

### Seizure terminating effects of closed-loop MS stimulation are mediated by GABAergic MS population

We next dissected contributions of the GABAergic, Glut and cholinergic MS neurons to the observed seizure terminating effects. Cell-type specific Cre-dependent expression of virally transduced ChR2 was established in three transgenic lines. As the VGAT- and CaMKIIα-Cre driver lines were newly developed using a CRISPR-Cas9 technology, we initially confirmed the cell-type specific ChR2 expression, and their proper electrophysiologic functionality (Dataset figures 5–9).

#### Closed-loop seizure rhythm-driven activation of MS GABAergic neurons terminates seizures of HPC-origin and prevents generalization

The MS of VGAT-Cre driver rats were injected with a Cre-dependent ChR2 expressing viral vector (Figure 4A). ChR2 signals were predominantly colocalized with glutamate decarboxylase (GAD) 65/67 immunoreactions but not with those of choline acetyltransferase (ChAT) (Figure 4B, Dataset figure 7). Some ChR2 signals were colocalized with glutamate reactions because glutamate is a precursor of GABA and MS GABAergic neurons use glutamate as local transmitters (38). MS illumination of the rats induced cFos expression in ChR2 expressing neurons (Figure 4C) and LFP deflections in the HPC and the EC but not in the Ctx (Figure S1E). Similarly to MS E-stim and hSyn∷ChR2 illumination, closed-loop seizure rhythm activation of MS GABAergic neurons significantly shortened HPC seizures compared with those in the interwoven non-illuminated trials (47.4 ± 21.9 s to 34.6 ± 18.3 s, *P* < 0.01, paired *t*-test, Figure 4D, E, G). The closed-loop GABAergic activation also shortened Ctx seizures and decreased Racine’s scale values (25.5 ± 7.6 s to 14.8 ± 16.6 s and 4.1 ± 1.3 to 2.7 ± 1.9 in Racine’s scale, *P* < 0.001 for both, paired *t*-test and Wilcoxon signed rank test, respectively; Figure 4E, G). Illumination and non-illumination trial pairs were classified into success and non-success trial pairs as before (*P* < 0.001, two-sample Kolmogorov-Smirnov test, Figure 4F) and this classification was consistently valid on Ctx seizures and motor seizures as well (*P* < 0.001 for all, three-way ANOVA, Figure 4G, I). On success trials, Ctx seizures were quite short and Racine’s scale values were less than three, which meant secondary generalization was effectively suppressed (< 7.25 s and < 2.25 in Racine’s scale, Figure 4I). Fractions of success trials were not significantly different over delays but there was a trend that larger delays (20–60 ms) were more effective than no delay (Figure 4H). These results suggest that GABAergic neurons mediate seizure terminating effects of closed-loop MS stimulation especially when stimulus was delivered with an appropriate delay. Open-loop activation of MS GABAergic neurons was also investigated (Figure S4Figure S3A, B). However, open-loop MS GABAergic activation did not shorten HPC or Ctx seizures and did not decrease Racine’s scale values (49.6 ± 21.8 s to 45.5 ± 27.4 s, 27.0 ± 16.1 s to 24.1 ± 16.4 s, and 4.3 ± 1.0 to 3.9 ± 1.2 in Racine’s scale, *P* = 0.38, 0.52, and 0.06, respectively; Figure S4C–F). These results suggest that seizure rhythm congruent activation is crucial for seizure terminating effects of MS GABAergic neurons as well.

#### Closed-loop seizure rhythm activation of MS Glut neurons does not effectively terminate seizures of HPC-origin

Roles of MS Glut neuronal populations were then investigated. The MS of CaMKIIα-Cre driver rats were injected with a Cre-dependent ChR2 expressing viral vector (Figure 5A). ChR2 signals were predominantly colocalized with glutamate immunoreactions but not with those of GAD67 or ChAT (Figure 5B, Dataset figure 8). MS illumination in the rats induced cFos expression in ChR2 expressing neurons (Figure 5C) and fast LFP deflections in the HPC and the EC but not in the Ctx (Figure S1E). Closed-loop seizure rhythm activation of MS Glut neurons did not significantly decrease duration of HPC and Ctx seizures, and Racine’s scale values in population data (53.2 ± 20.9 s to 50.6 ± 17.2 s, 27.8 ± 15.4 s to 26.4 ± 15.0 s, and 4.5 ± 1.1 to 4.2 ± 1.5, *P* = 0.37, 0.46, and 0.10, respectively; Figure 5D, G). Distribution of MIs of HPC seizures has its mode around zero; however, the distribution has a longer tail toward −1, which meant that there were success trial pairs (Figure 5F). Thus, the threshold was introduced to classify the MI distribution into success and non-success trial pairs. This classification was consistently valid on Ctx seizures and Racine’s scale as well (*P* < 0.001, Figure 5F, H). Although still low, the fraction of the success trials with none-delayed illumination in CaMKIIα∷ChR2 rats tends to be higher than those with delays (0.200, 0.039, 0.067, and 0.067 for 0, 20, 40, and 60 ms delay, respectively; Figure 5E, I). This result on delay was complementary with those with VGAT∷ChR2 rats; activation of MS GABAergic population was more effective with 20‒60 ms delay after each seizure wave detection (Figure 5I). The sum of fractions of success trial pairs from CaMKIIα∷ChR2 and VGAT∷ChR2 experiments is similar to that of MS E-stim experiments (0.422, 0.333, 0.361, and 0.537 for CaMKIIα + VGAT and 0.539, 0.429, 0.429, and 0.436 for E-stim, respectively; Figure 5I).

**Figure 5.**
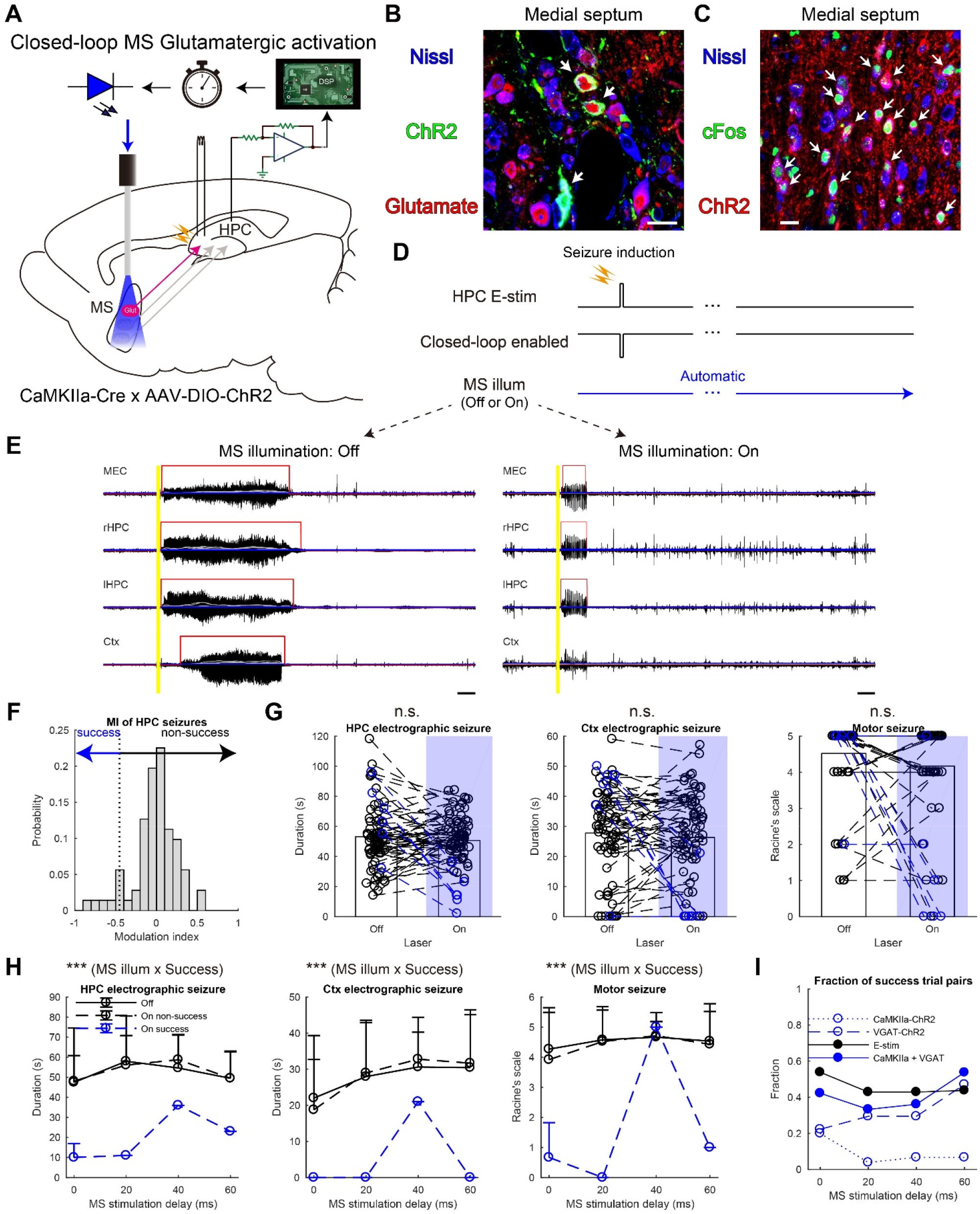
Closed-loop seizure rhythm-driven activation of MS Glut neurons does not effectively terminate seizures of HPC-origin. (**A**) Schema of the experiment. (**B**) ChR2 signals were colocalized with Glut neurons. (**C)**cFos expression was colocalized with ChR2 expressing neurons after illumination. (**D**) Closed-loop seizure rhythm intervention. (**E**) Representative seizure waves w/wo the MS illumination in CaMKIIα∷ChR2 rats. (**F**) Distribution of MIs with the MS illumination in CaMKIIα∷ChR2 rats. Each trial pair was clustered into success or non-success group by splitting at the global threshold. (**G**) Duration of HPC and Ctx seizures, and motor seizures with the clusters defined in (F). (**H**) MS illumination delay resolved representation of the data shown in (**G**). (**I**) Fractions of success trial pairs as a function of delays in rats with MS CaMKIIα∷ChR2 illumination. Fractions of success trial pairs in rats with MS VGAT∷ChR2 illumination and MS E-stim are re-plotted from Figures 4H and 3G, respectively. The summation plot of CaMKIIα∷ChR2 and VGAT∷ChR2 data (CaMKIIα+ VGAT) is similar to that of E-stim data. n = 142 trials from three CaMKIIα∷ChR2 rats. Other conventions are the same as Figure 4. See also Figure S5.

Open-loop MS Glut activation was also investigated (Figure S5A, B). However, the open-loop MS Glut activation did not shorten HPC or Ctx seizures and did not decrease Racine’s scale values at all (53.2 ± 21.0 s to 50.6 ± 17.2 s, 27.8 ± 15.4 s to 26.4 ± 15.0 s, and 4.5 ± 1.1 to 4.2 ± 1.5 in Racine’s scale, *P* = 0.08, 0.85, and 0.26, respectively; Figure S5C–F).

Together, these results suggest that activation of the MS Glut neuronal population alone does not effectively terminate HPC-origin seizures. Precisely timed activation of MS Glut neurons might have a relatively minor contribution to the seizure terminating effects of seizure rhythm MS activation with distinct phase preference from that of the activation of GABAergic population.

#### Neither closed-loop nor open-loop activation of MS cholinergic neurons alleviates seizures of HPC-origin

The MS of ChAT-Cre driver rats were injected with a Cre-dependent ChR2 expressing viral vector (Figure S6A, F). ChR2 signals were predominantly colocalized with ChAT immunoreactions but not with those of GAD65/67 (Figure 6F, Dataset figure 9). Some ChR2 signals were colocalized with glutamate reactions as well, which is presumably because glutamate is local transmitter also in MS cholinergic neurons (38). MS illumination in the rats did not induce deflection of HPC LFPs (Figure S1E) but induced cFos expression in ChR2 expressing neurons (Figure 6G, Dataset figure 9). In contrast to GABAergic and Glut activation, closed-loop seizure rhythm activation of MS cholinergic neurons did not alleviate HPC, Ctx, or motor seizures (55.0 ± 17.8 s to 51.2 ± 21.4 s, 39.2 ± 17.8 s to 36.0 ± 19.5 s, and 4.8 ± 0.6 to 4.8 ± 0.8 in Racine’s scale, *P* = 0.28, 0.22, and 0.83, respectively; Figure S6B, C, E). MI distribution was not skewed (*P* = 0.51, two-sample Kolmogorov-Smirnov test, Figure S6D).

**Figure 6.**
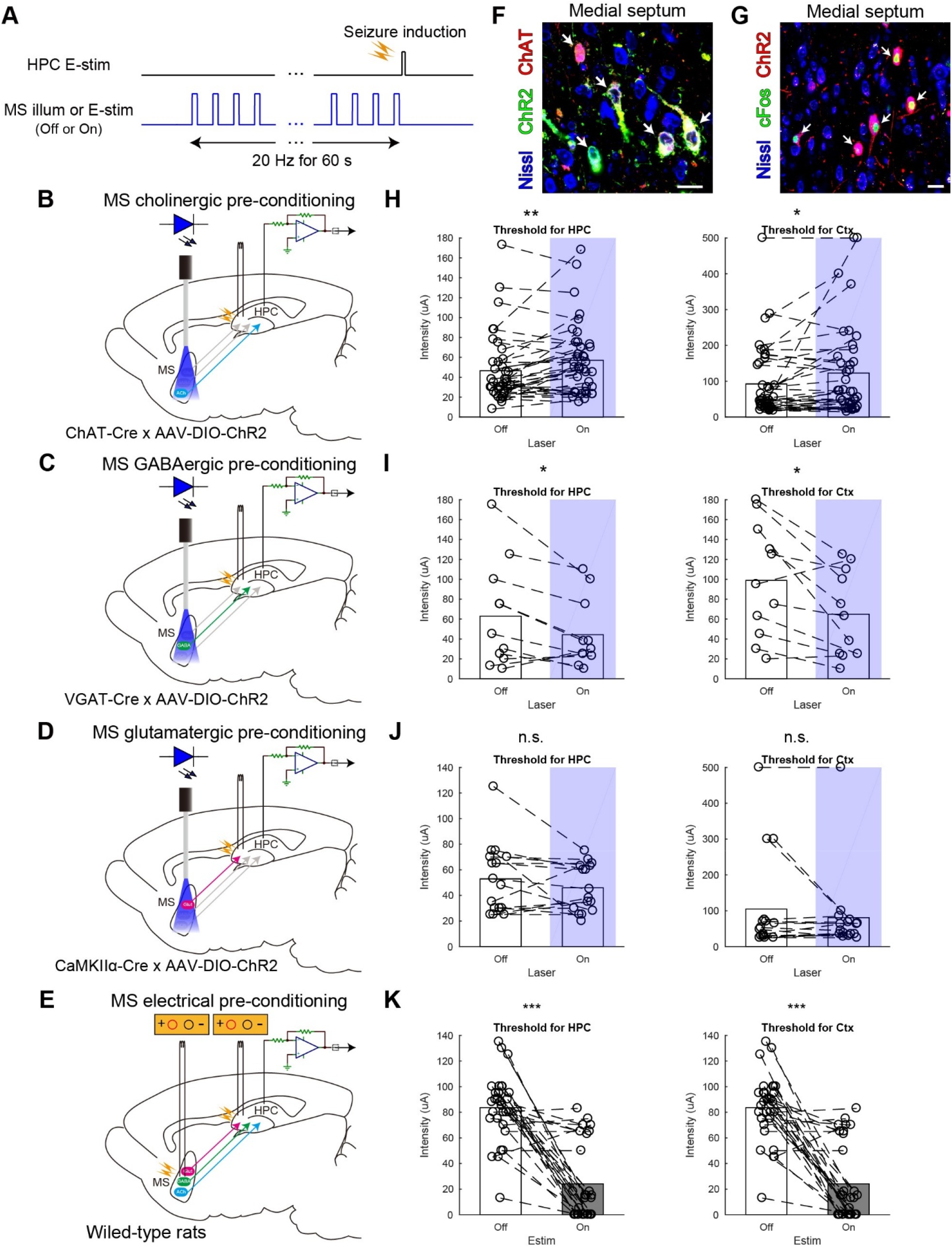
MS cholinergic pre-conditioning decreases seizure susceptibility whereas MS E-stim pre-conditioning at seizure-like-frequency increases it. (**A**) Pre-conditioning procedure. MS neurons were optogenetically or electrically activated at 20 Hz for 60 s before seizure induction. (**B**–**E**) Schemata of the MS optogenetic pre-conditioning. MS cholinergic, GABAergic, or Glut neurons were selectively activated, respectively. (**E**) Schema of the MS electrical pre-conditioning. (**F**) ChR2 signals were colocalized with cholinergic neurons in ChAT-Cre driver rats injected with a Cre-dependent ChR2 expressing viral vector. (**G)**cFos expression was colocalized with ChR2 expressing neurons in the rats after illumination. (**H**–**J**) Stimulus intensity thresholds for inducing HPC and Ctx seizures w/wo pre-conditioning in MS cholinergic, GABAergic, or Glut neurons, respectively. (**K**) Stimulus intensity thresholds for inducing HPC and Ctx seizures w/wo electrical pre-conditioning. n = 78 trials from four rats for (H), 22 trials from three rats for (I), 32 trials from two rats for (J), and 60 trials from two rats for (K), respectively. Values are represented as columns for means and a marker for each trial. Statistical significance was tested by paired *t*-test. n.s., not significant; \**P* < 0.05; \*\**P* < 0.01; \*\*\**P* < 0.001. See also Figure S6.

Open-loop activation of the MS cholinergic neurons was then examined but it again did not shorten HPC, Ctx, or motor seizures (51.8 ± 30.7 s to 53.5 ± 29.6 s, 29.6 ± 20.6 s to 32.5 ± 22.5 s, and 4.1 ± 1.6 to 3.9 ± 1.6 in Racine’s scale, *P* = 0.79, 0.50, and 0.58, respectively; Figure S6F, G, H, J). MI distribution was not skewed (*P* = 0.39, two-sample Kolmogorov-Smirnov test, Figure S6I). These results suggest that phasic activation of MS cholinergic neurons during ictal periods is not effective to alleviate seizures of HPC-origin.

### MS cholinergic pre-conditioning decreases seizure susceptibility whereas MS E-stim pre-conditioning at seizure-like-frequencies increases it

Finally, driven by our idea of preventing seizure development through prediction of ictal activity, we investigated whether pre-activation of any septal neuronal populations at seizure-like frequency decreases seizure susceptibility. We found earlier that MS cholinergic activation can decrease high-frequency HPC oscillations (17). Thus, we first activated MS cholinergic neurons in kindled rats at 20 Hz for 60 s before seizure induction (Figure 6A, B). With repeating this pre-conditioning, stimulus intensity on the HPC commissure was incremented over trial and trial to determine threshold intensities to trigger HPC and Ctx seizures. Cholinergic pre-conditioning increased thresholds for HPC and Ctx seizures (46.6 ± 34.1 μA to 57.1 ± 35.7 μA and 92.7 ± 99.5 μA to 122.7 ± 129.4 μA, respectively; *P* < 0.05 for both, paired *t*-test, Figure 6H). Five-second-long pre-conditioning was not enough to decrease the seizure susceptibility (data not shown). In contrast to cholinergic signaling, pre-activation of GABAergic neurons decreased thresholds for HPC and Ctx seizures (63.0 ± 53.0 μA to 44.3 ± 34.6 μA and 98.9 ± 56.9 μA to 64.9 ± 43.2 μA, respectively; *P* < 0.05 for both, paired *t*-test, Figure 6C, I). Pre-activation of Glut neurons did not cause any change in the thresholds (52.9 ± 27.4 μA to 45.9 ± 18.4 μA and 105.1 ± 137.6 μA to 80.6 ± 114.0 μA, *P* = 0.21 and 0.21 for thresholds for HPC and Ctx seizures, respectively; paired *t*-test, Figure 6D, J). To determine net effects of simultaneous pre-activation of these three neuronal populations, electrical pre-conditioning of the MS was next conducted (Figure 6E). MS E-stim at 20 Hz for 60 s before seizure induction significantly decreased thresholds for HPC and Ctx seizures (83.6 ± 25.7 μA to 24.1 ± 30.2 μA for both, note that thresholds for HPC and Ctx seizures were the same in all trials, *P* < 0.001, paired *t*-test, Figure 6K). Notably, 20 Hz MS E-stim itself occasionally induced seizures during the pre-conditioning exposure. These results suggest that selective activation of MS cholinergic neurons during pre-ictal periods could prevent seizure development. However, this seizure preventing effect is cell-type specific because the net effect of MS E-stim at 20 Hz (seizure like frequency) is pro-seizure. Therefore, MS E-stim at seizure-like-frequency should not be delivered during pre-ictal periods; it should be delivered rather during ictal periods in a closed-loop manner, precisely aligned to internal seizure patterns.

## Discussion

We found that properly performed MS stimulation can terminate epileptic seizures of HPC-origin and can prevent their generalization. Notably, there is a dramatic difference between outcomes of closed-loop and open-loop interventions; closed-loop seizure rhythm-driven MS stimulation significantly shortens HPC seizures whereas open-loop stimulation does not shorten or even prolongs them. Considering significant correlations between modulation of HPC seizures and those of Ctx and motor seizures (Figure S7), we claim bidirectional roles of the MS on TLE: anti- and pro-seizure effects (Figure 7). Our cell-type specific optogenetic experiments revealed that the MS GABAergic neuronal population mediated mainly the anti-seizure effects of MS.

**Figure 7.**
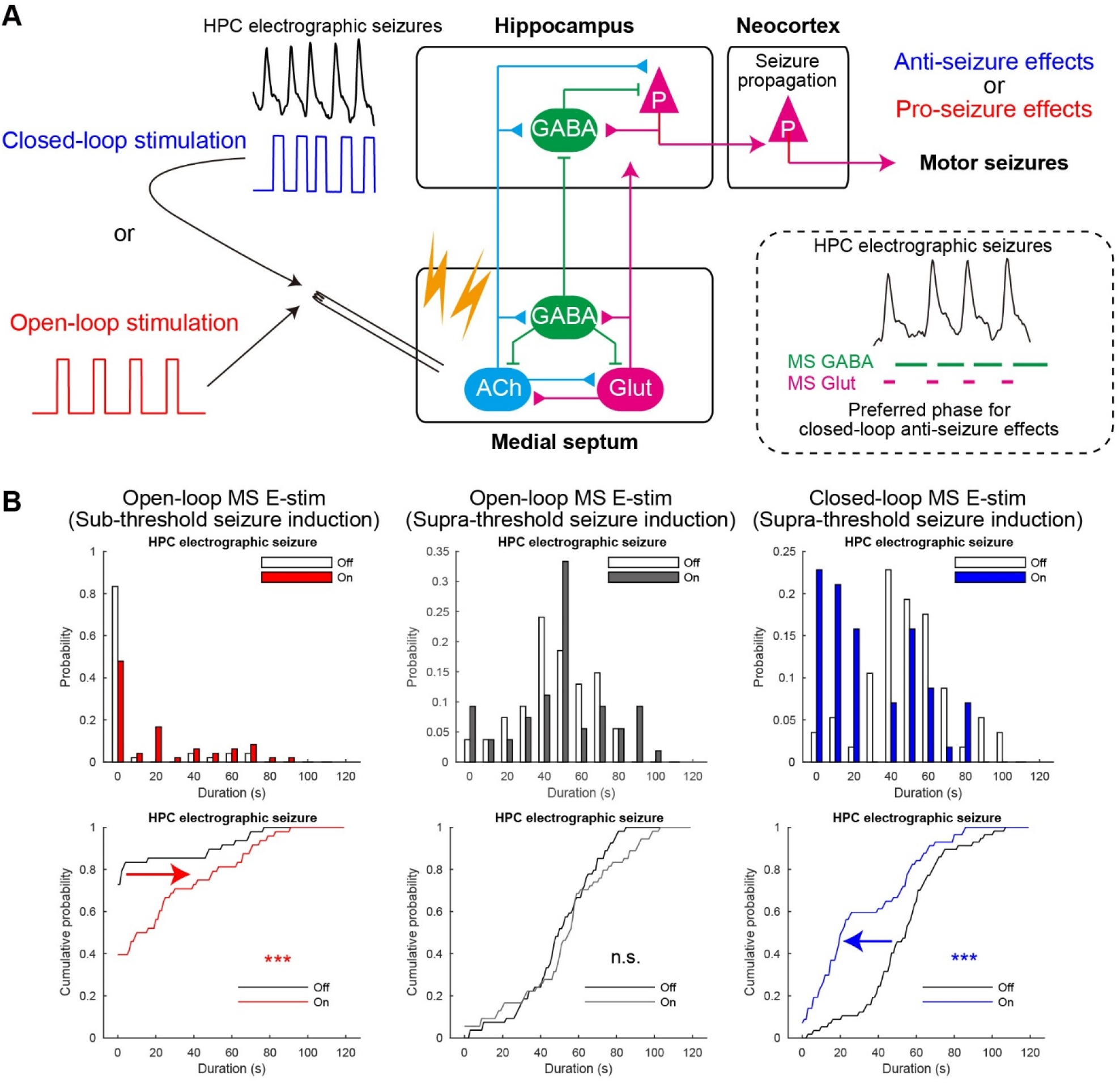
Summary. (**A**) Closed-loop seizure rhythm-driven MS E-stim is effective to terminate seizures of HPC-origin and successfully suppresses secondary generalization. Alternating activation of MS GABAergic and Glut neurons may underlie the seizure terminating effect. In contrast, open-loop responsive MS E-stim does not alter or rather promotes HPC seizures. (**B**) Distributions and cumulative probabilities of HPC seizure duration w/wo MS E-stims. Data are from Figures 2 and 3. The red and blue arrows indicate significant pro- and anti-seizure effects. Statistical significance was tested by two-sample Kolmogorov-Smirnov test. n.s., not significant; \*\*\**P* < 0.001. See also Figure S7.

### MS as a target of DBS in refractory TLE: dual roles in anti-seizure effects

The MS has dual roles regarding its anti-seizure effects: on-demand seizure terminating effects and seizure susceptibility reducing effects. Regarding the former, such powerful on-demand seizure terminating effects on already initiated episodes have not been reported yet with other proxy interventions in refractory TLE. E-stim of the anterior nucleus of the thalamus (ANT) (12), the cortex (21), the vagus nerve (39) achieved approximately 40–50% reduction of seizures occurrence in TLE patients. However, the remaining 50–60% of the episodes, which may become secondary generalized, remain to be treated. The powerful seizure terminating effect of the MS presumably stems from driving its direct reciprocal connections to the bilateral HPCs and the ECs in a temporally precise manner (26, 40). Regarding the other role, previous studies have shown that open-loop theta (4–8 Hz) MS E-stim reduces seizure susceptibility in animal models (41, 42). This theta rhythm MS stimulation has been thought to restore disrupted theta oscillations in the epileptic HPC via presumably cholinergic mechanisms (43, 44). The dual roles of the MS for anti-seizure effects make it valuable as a target of DBS for refractory TLE.

### Advantages of MS stimulation in TLE

We point out three additional advantages of MS stimulation. First, because the MS is located at the midline, there is no need to insert multiple electrodes for bilateral neuromodulation. This is advantageous compared to stimulation of the ANT, the HPC, and the Ctx (8), as for example the ANTs are bilaterally located and each of them is indirectly connected only to the ipsilateral HPC via the Papez circuit (45). Second, electrical stimulation can be employed for clinical practice: no need of optogenetics. Although cell-type and phase specific seizure terminating effects of the MS were revealed, only a seizure rhythm congruent stimulus delivery is practically required for the seizure terminating effects. Third, both seizure rhythm and theta rhythm stimulation can be delivered in an on-demand manner. This time-targeting (responsive) nature is a significant advantage over pharmaceutical treatments and traditional continuous DBS (21). Practically, continuous MS stimulation should be avoided so as not to disrupt cognitive functions supported by natural oscillations in the septohippocampal axis (28, 46) and to prevent kindling effects. Repeated stimulus of the limbic system including the MS may make the circuit epileptic (47). Adaptive, but continuous DBS with real-time modulation of stimulus amplitude (48) may not be suitable for the MS because even sub-threshold stimuli may induce kindling effects there. Responsive (on-demand) stimulus delivery to the MS will overcome the risk of kindling effects with the continuous DBS. Together, responsive seizure rhythm and theta rhythm MS stimulation for ictal and predicted pre-ictal periods would be beneficial to achieve successful control on TLE.

### Possible mechanisms of seizure terminating effects of closed-loop MS stimulation

#### Network mechanisms

Three network mechanisms could underlie the seizure terminating effects. The first one is on network stability. Seizure networks have multi-stable states as interictal, pre-ictal, and ictal states (49, 50). Huge perturbation on network dynamics during ictal states would cause state transitions to another state. Each MS stimulation may have only a small impact. However, if many stimuli are time-locked to a certain phase of seizure oscillations, either resonating with (in-phase) or occluding it (counter-phase), their impacts would be temporally summed. The network dynamics could be then pushed away from the ictal state as a swing collapse. For example, counter-phase activation of MS or HPC projection neurons could occlude seizure oscillations with their refractory periods. In addition, antidromic spikes of MS targeting HPC neurons presumably induce collisions on their axons and occlude seizure oscillations as well. Even excess in-phase positive feedback may disrupt stability of seizure states like a howling phenomenon (51). Open-loop stimulation would not cause such resonance and result in small impacts. The second network mechanism is a competition between different oscillation frequencies. Newly added oscillations influence pre-existing oscillations (30). Neuronal networks have their own resonating frequency and because closed-loop seizure rhythm stimulation would be a similar oscillation pattern to the original oscillation, the synthetized wave would be harmonics of the original one, which could reduce the original oscillation. The third one is a bottle neck effect. Seizures of HPC-origin reverberate and are amplified in the HPC-HC and the HPC-MS loops. To be secondary generalized, the EC is then passed through, which is the bottle neck structure. MS GABAergic neurons project not only to the HPC but also the EC in a very diffuse manner. The projection is primarily targeting interneurons in the EC as well as in the HPC (52). EC and HPC interneurons have wide-spread oscillation gating effects there: hub-like effect (53, 54). These proxy and hub-like effects could be then synergistically amplified by temporal summation on oscillation cycles to gate the EC.

#### Cellular mechanisms

The blue light illumination of MS cell populations in VGAT∷ChR2, CaMKIIα∷ChR2, and ChAT∷ChR2 rats induced very distinct spatiotemporal patterns of LFP deflection in the HPC and EC (Figure S1) and seizure terminating effects (Figures 4–6, S4–6). Our failure to terminate seizures in CaMKIIα∷ChR2 animals confirm that the observed effects of our optogenetic experiments would not be artifacts of illumination or heating itself, which would happen even without any opsins (55), but are cell-type specific neuronal responses.

MS GABAergic neurons projects to the HPC and the EC and terminate there on mainly GABAergic neurons (56). Phasic activation of MS GABAergic neurons during trough phase (20–60 ms from peak) would induce rebound firing of HPC GABAergic neurons. The rebound activation then introduces powerful perisomatic inhibition on HPC pyramidal neurons around the next peak of seizure oscillations, when population burst of pyramidal neurons synchronously occurs (57). This time-targeted inhibitory effects on pyramidal neurons can be temporally summed by the phase-locked nature and spatially amplified by wide-spread axonal innervation of MS GABAergic neurons to the HPC and the EC (52). Because HPC GABAergic neurons are hypofunctional during TLE, which leads to epileptiform discharges of pyramidal neurons, this time-targeted phasic activation of HPC GABAergic neurons via the MS would be essential for terminating epileptic discharges in the HPC.

Although relatively weekly, MS Glut neurons might also mediate the seizure terminating effects. Interestingly, Glut and GABAergic interventions had complemental preferred phases to achieve their best seizure terminating effects (Figure 5I). Because MS Glut neurons have intense intra-septal innervation on GABAergic neurons and their direct innervation to the HPC and EC is minor, their seizure terminating effects would be mediated through MS GABAergic neurons. The approximately 20 ms temporal shift between MS Glut and GABAergic activation can stem from synaptic delays between them, and the time required to cumulate a significant gross glutamatergic effect on the GABAergic population. The tri-synaptic pathway of MS Glut neurons -> MS GABAergic neurons -> HPC GABAergic neurons -> HPC pyramidal neurons can deteriorate temporal summation for the seizure terminating effects via cumulating synaptic jitters and varying transmission efficacy. This can be a reason why MS Glut activation has weaker seizure terminating effects compared to those of MS GABAergic neurons.

The on-demand seizure terminating effect is not grossly influenced by the septohippocampal cholinergic signalling because this latter is mainly mediated by slow metabotropic receptors (34), which cannot follow rhythmicity of the seizure oscillations. Additional anti-seizure effects by HPC cholinergic tone during ictal periods could not be observed. In contrast, activation of MS cholinergic neurons in the pre-ictal periods reduced seizure susceptibility (Figure 6). This is consistent with our report that activation of MS cholinergic neurons decreased occurrence of high-frequency episodes in the HPC (17). Oriens-lacunosum moleculare cells may mediate these effects (58).

Our experiments revealed the seizure terminating effect of closed-loop intervention in a provoked seizure model. The temporal organization of the stimulation may need to be adjusted if using other chronic spontaneous seizure models, since their underlying cellular and network mechanisms may be different. The feasibility of the MS induced seizure termination in other seizure models shall be investigated in the future.

### Possible future clinical application

MS E-stim has been suggested for clinical studies after the finding that theta rhythm MS stimulation reduces seizure susceptibility (59, 60). Together with the previous findings on open-loop stimulation of the MS (41, 42), our current results indicate that a single electrode in the MS can be used for dual purposes: seizure termination by closed-loop seizure rhythm-driven stimulation during ictal periods and seizure prevention by open-loop theta rhythm stimulation during pre-ictal seizure initiation periods. Development of reliable seizure prediction and detection algorithms for spontaneous seizures in humans would be a future challenge and is out of the scope of this work, as we only employed a provoked seizure models in rats. Stability of closed-loop stimulation should be optimized by decreasing particularly false positive detections of stimulus artifacts and other ambient noises. Longitudinal, patient specific fine-tunings of detection and stimulus algorithms would be important to ensure long-term effectiveness and safeness of MS E-stims. Intelligent DBS approaches utilizing deep learning technology may be effective for real-time classification of brain states to deliver proper stimulus patterns in an on-demand manner (61).

In the present study, we showed that closed-loop seizure rhythm-driven MS E-stim successfully terminated HPC seizures preventing its generalization, and provided a concept for a novel therapeutic approach of TLE directly translatable to clinical practice. The concept that precisely time-targeted proxy stimulation can produce massive impacts on pathological oscillations could be extrapolated in the future to regulate other oscillopathies including movement or mental disorders (62).

## Author Contributions

Conceptualization, A.B.; Methodology, Y.T., G.K., and A.B.; Software, Y.T. and G.K., Validation, Y.T. and A.B.; Formal Analysis, Y.T. and T.F.; Investigation, Y.T., M.H., and L.P.; Resources, Y.T., G.K., and A.B.; Data Curation, Y.T.; Writing – Original Draft, Y.T., Writing – Review & Editing, A.B.; Visualization, Y.T.; Supervision, A.B.; Project Administration, Y.T. and A.B.; Funding Acquisition; Y.T., M.H., G.K., and A.B.

## Acknowledgments

We thank Ms. Mari Takeuchi, Dr. Qun. Li and Dr. Masahiro Ohsawa for technical assistance, Dr. Karl Deisseroth for ChR2-expressing viral vectors and ChAT-Cre rats, Dr. Péter Hegyi for providing access to confocal microscopy. This work was supported by KAKENHI (18KK0236, 19H03550, 19H05224), the Uehara Memorial Fund, the Kanae Foundation for the Promotion of Medical Science, and Life Science Foundation of Japan to Y.T., the Szeged Scientists Academy under the sponsorship of the Ministry of Human Capacities, Hungary (EMMI:13725-2/2018/INTFIN) to M.H., the UNKP-18-3 of the Ministry of Human Capacities, Hungary to G.K., EU-FP7-ERC-2013-Starting grant (No. 337075), EU H2020 No. 739593, Momentum program I and II of the Hungarian Academy of Sciences, EFOP-3.6.1-16-2016-00008 - National Research, Development and Innovation Office, Hungary, and 20391-3/2018/FEKUSTRAT of the Ministry of Human Capacities, Hungary to A.B.

## Materials and Methods

### CONTACT FOR REAGENT AND RESOURCE SHARING

Further information and requests for resources and reagents should be directed to and will be fulfilled by the Lead Contact, Antal Berényi.

### EXPERIMENTAL MODEL AND SUBJECT DETAILS

#### Rat models

All experiments were performed in accordance with European Union guidelines (2003/65/CE) and the National Institutes of Health Guidelines for the Care and Use of Animals for Experimental Procedures. The experimental protocols were approved by the Ethical Committee for Animal Research at the Albert Szent-Györgyi Medical and Pharmaceutical Center of the University of Szeged (XIV/218/2016). In total 50 adult Long-Evans rats (eight wild type and 42 transgenic, 3–6 months old, 300–640 g) were used in the present study. Both sexes of rats were used. VGAT-Cre rats (LE-*Slc32a1*^tm1(cre)*Sage*^; RGD Cat# 12905033; RRID:RGD_12905033) and CaMKIIα-Cre rats (LE-*Camk2a^tm1(IRES-cre)Sage^*, RGD Cat# 12905032; RRID:RGD_12905032) were purchased and licensed from Horizon Discovery (Cambridge, UK). ChAT-Cre rats: LE-Tg(ChAT-Cre)5.1Deis, RGD Cat# 10401204; RRID:RGD_10401204) were given by Dr. Deisseroth (63). Rats were fed a commercial diet and water *ad libitum* with controlled temperature and lighting (12/12 h light/dark cycle) and housed as groups. Rats were either bred from in house colonies or purchased from Charles River (Sulzfeld, Germany). No animals were excluded from analysis.

### METHOD DETAILS

#### Genotyping

ChAT-Cre rats were maintained as male heterozygotes and genotyped by a standard PCR procedure with the following primers: CW-Cre2, 5’-ACC TGA TGG ACA TGT TCA GGG ATC G-3’ and CW-Cre3, 5’-TCC GGT TAT TCA ACT TGC ACC ATG C-3’, producing 108-bp fragments from the *cre* allele (64). Other Cre driver lines were maintained as homozygotes of both sexes after initial genotyping using the manufacturer’s PCR protocols. No randomization or blinding was employed for experiments below.

#### Stereotaxic viral vector injections

For ChR2 expression, AAV5-hSyn-hChR2(H134R)-mCherry or AAV5-EF1α-DIO-hChR2(H134R)-mCherry (UNC Vector Core, Chapel Hill, NC, USA) was injected into the MS of wild-type or Cre-driver animals using a standard intracranial virus vector injection technique, respectively. Briefly, animals were anesthetized with 1–3% isoflurane and then mounted on a stereotaxic apparatus. Atropine (0.1 mg/kg, s.c.) was administered immediately after the anesthesia induction. Stages of anesthesia were maintained by confirming the lack of nociceptive reflex. The rectal temperature was maintained at 36–37°C with a DC temperature controller (TMP-5b; Supertech, Pécs, Hungary). The head position was adjusted so that the bregma and lambda were at the same level. Small incision on the skin was then made for craniotomy after subcutaneous lidocaine injections. A craniotomy centered at 1.0 mm leftward from the midline and 0.5 mm anterior from the bregma was drilled and a small incision on the dura mater was made. A glass capillary (tip 10–20 μm) filled with an AAV vector without dilution (qPCR titer: 3–5×10^12 vg/ml) was installed with an auto-nanoliter injector (Nanoject II; Drummond Scientific, Broomall, PA, USA) and the injector was 9.5 degree leftward tilled from the sagittal plane to avoid the sagittal sinus. Three penetrations were made from the following stereotaxic coordinates: 0.2, 0.5, and 0.8 mm anterior from the bregma and 1.0 mm leftward from the midline. Along each penetration, 0.3 μl viral solution was ejected at 5.8, 6.1, and 6.4 mm from the pia at a rate of 0.9 nl/s. The capillary tip was kept there for at least five minutes after each ejection and then gently retracted. The cranial window was covered with a hemostatic gel, and the skin wound was sutured. The injected animals were housed at least four weeks before optogenetic intervention.

#### Electrophysiology

To monitor electrographic seizures, intracranial LFP recordings were made from freely-moving animals. Intra-MS Electrophysiological correlates of optogenetic stimulation were confirmed in anesthetized animals (Dataset figure 5)

##### Construction of recording electrodes

Tripolar tungsten wire electrodes were prepared as previously reported (65). Briefly, we glued three 50 μm diameter HML-insulated tungsten wires (Tungsten 99.95%; California Fine Wire, Grover Beach, CA, USA) into a 25G stainless-steel tube. Tips of the tungsten wires were spaced at a 0.4 mm axial distance. Impedance of each wire was 30–90 kΩ at 1 kHz.

##### Chronic implantation surgery

Each animal was implanted with 10 tripolar electrodes under stereotaxic guidance as previously described (65). In total 30 recording sites were distributed to the bilateral dorsal hippocampus (rHPC, lHPC), right medial entorhinal cortex (MEC), and right somatomotor cortex (S1/M1). The stereotaxic coordinate of recording sites was as follows: S1/M1 (AP: 1.0 and 2.5 mm anterior from the bregma; ML: 3.0 mm; DV: 1.2, 1.6, and 2.0 mm from the dura), HPC (AP: 3.5, 4.5, and 5.5 mm posterior from the bregma and ML: 2.0, 3.0, and 4.0 mm, respectively; DV: 2.6, 2.9, and 3.3 mm from the dura), MEC (AP: penetrated from the 0.1 mm anterior from the transverse sinus at a 20° caudally tilted angle from the coronal plane; ML, 3.5 and 4.5 mm; Distance: 3.2, 3.6, and 4.0 mm from the dura). The dura matter was not removed but penetrated with the tripolar electrodes. After insertion, the electrodes were physically fixed with a dental cement (Unifast Trad; GC, Tokyo, Japan). Two stainless-steel machine screws were installed in the skull above the cerebellum as reference and ground electrodes. A bipolar stimulus electrode was placed targeting the HPC commissure for kindling and seizure induction during the experiments (see below). Either an optical cannula or a bipolar stimulus electrode was inserted into the MS as well. Finally, an on-head Faraday cage made of copper mesh connected to the ground electrode was cemented on the skull to prevent environmental noise and physically protect the implanted components. The implanted animals were housed in individual cages.

##### Data acquisition from freely-moving animals

Daily LFP recordings were performed in the animals’ own home cages. The tungsten wire electrodes were connected to a signal multiplexing headstages (HS3_v1.3, Amplipex, Szeged, Hungary) attached to a thin and light cable (36 AWG Soveron served Litz wire; Kerrigan-Lewis, Alpha wire, Elizabeth, NJ, USA) pending from a trolley system on the room ceiling, which allowed free moving of the animals. To avoid twisting and over-tension of the recording cables and an optical fiber for optogenetic stimulation, a bore-through electrical commutator (VSR-TC-15-12; Victory-Way Electronic, Shenzhen, China) and an optical rotary joint (RJPSF2; Thorlabs, Newton, NJ, USA) were integrated into the recording cables. The multiplexed signals were acquired at 500 Hz per channel for closed-loop intervention experiments (23) and at 20 kHz per channel for open-loop intervention experiments and other recordings (KJE-1001, Amplipex) (66).

##### Post-mortem localization of the recording sites

After data acquisition, animals were deeply anesthetized with 1.5 g/kg urethane (i.p.). One or two recording sites of each tripolar electrode were lesioned with 100 μA anodal direct current for 10 s via tips. The animals were transcardially perfused and their brains were sectioned as described below. The sections were stained with 1 μg/ml 4′,6-Diamidino-2-phenylindole dihydrochloride in distilled water (D8417; Sigma-Aldrich, St. Louis, MO, USA), cover-slipped, and observed with a confocal microscopy (Figure S1).

#### Hippocampal electrical kindling

##### Kindling procedure

A bipolar stimulus electrode for kindling and seizure induction was prepared using the same tungsten wire as tripolar recording electrodes. Tips of the bipolar stimulus electrodes were axially spaced at 0.5 mm distance and insulation 0.4–0.5 mm around the tips were stripped to decrease their impedance to 10–20 kΩ at 1 kHz. The stimulus electrode was implanted on the HPC commissure during the chronic implantation surgery for recording electrodes in the following stereotaxic coordinate: AP, 0.84 mm posterior from the bregma; ML, penetrated from 1.5 mm rightward from the midline at a 18° rightward tilted angle from the parasagittal plane; Distance, 4.5 mm from the dura. The kindling electrode was fixed with dental cement. After recovery from the implantation surgery, the HPC was daily stimulated at sub-convulsion intensity through the kindling electrode. Each kindling stimulation consisted of 120 × 0.5 ms positive-0.5 ms negative bipolar rectangular pulses at 62.5 Hz (36) generated via a programable isolator in current control mode (STG4008; Multi Channel Systems, Reutlingen, Germany). Stimulus intensity was determined as the minimum intensity that induced after-discharge (population spikes) in HPC LFPs, which was typically ±20–200 μA. The kindling stimulation was conducted six times per day for 10 days and each kindling stimulation was delivered with at least 30 min inter-stimulus intervals to develop secondary generalized seizures in response to the kindling stimulation (Dataset figure 1, see Table S2).

##### Behavioral monitoring

Behavior of animals were continuously monitored during electrophysiological recordings with a webcam (LifeCam Studio; Microsoft, Redmond, WA, USA) in VGA resolution and in 30 fps rate in a synchronized manner to the acquired neuronal data. Severity of motor seizures was evaluated on-line and off-line according to the Racine scale: 1, Mouth and facial movements; 2, Head nodding; 3, Forelimb clonus; 4, Rearing; 5, Rearing and falling (47). Number of wet-dog shakes on each stimulation trial was also counted, which negatively correlated with a fully kindled state (67).

#### Optogenetics

##### Construction of optical cannulas

One side of 0.39 NA, Ø200 μm core multimode optical fiber (FT200EMT, Thorlabs) was terminated with a stainless-steel ferrule (SF230, Thorlabs). The buffer on the other side were 10 mm-removed with a fiber stripping tool (T12S21, Thorlabs) and the TECS cladding was then removed using an acetone-soaked cotton bud. After that, the exposed silica core was etched with hydrofluoric acid to form its tip as pencil-like shape, which facilitates penetration and increases illumination volume. Current-output power relationships were examined on every optical cannula with a photodiode power sensor (S130C, Thorlabs) and a power meter (PM200, Thorlabs). The typical maximal power at the tip was set to 30–40 mW.

##### Construction of optoprobes

A multimode Ø50 μm optical fiber (FG050LGA, Thorlabs) was attached to each silicon probe based on a method previously described (68). Briefly, one side of the fiber was terminated with a stainless-steel LC ferrule (SFLC127, Thorlabs) in accordance with a manufacturer’s guide (FN96A, Thorlabs). The acrylate coating of the other side was removed at 1 cm-long, fluorine-doped cladding was then thinned with hydrofluoric acid while the silica core was kept intact. After that, each etched optical fiber was mounted on a single shank 32 channel silicon probe (A1×32-Poly2-10mm-50s-177-H32_21mm; NeuroNexus, Ann Arbor, MI, U.S.A.) with a UV-curing optical adhesive (NOA61, Thorlabs). Current-output power relationships were examined on every optoprobe with a photodiode power sensor (S130C, Thorlabs) and a power meter (PM200, Thorlabs). The typical maximal power at the tip was set to 1–2 mW.

##### Light source and its control

A 450 nm laser diode (PL450B; Osram, Munich, Germany) was driven via a current controller with analog modulation (LDC205C; Thorlabs), and the emission beam was collimated with an aspheric lens. The collimated ray was divided and directed into two channel fiber ports, each of which is connected to a rat via multimode patch cables and a rotary joint. The laser power on each fiber port can be controlled between 0–100 mW with a polarizing beam splitter, a half-wave plate, analog modulation, and manual shutters (Dataset figure 4). Laser power was stabilized with a self-made feedback control system consisted of a photo detector (PDA36A-EC, Thorlabs) and a microcomputer (STM32L152RE; STMicroelectronics, Geneva, Switzerland). External command voltage was provided from a data acquisition board (USB-6212; National Instruments, Austin, TX, USA) or a pulse stimulator (Master-9; A.M.P.I., Jerusalem, Israel).

##### Unit recordings of MS neurons and their optogenetic modulation

Animals with ChR2 expression in the MS were anesthetized with 1.5 g/kg urethane (i.p.) and 0.1–0.3 mg/kg atropine (s.c.) and mounted on a stereotaxic instrument. Depth of anesthesia was maintained by confirming the lack of nociceptive reflex. The rectal temperature was maintained at 36–37°C with a DC temperature controller. The head position was adjusted so that the bregma and lambda were at the same level. A small cranial window (approx. 1 mm diameter) was made at 0.5 mm anterior from the bregma and 1 mm leftward from the midline. An optoprobe was inserted at an angle 9.5° leftward tilted from the parasagittal plane and gently advanced toward the MS. The electrode was kept and stabilized at several sites in the MS (5700–7000 μm from the pia) for recordings. At each recording site, pulse and sine wave illuminations paired with non-illuminating epochs were applied to find optogenetically modulated units. The parameters for pulse illumination were as follows: duration: 10 ms; amplitude: typically 1 mW; frequency: 1, 5, 10, 20, and 40 Hz; number of trials: 20; minimum number of cycles in each trial: 20; minimum trial duration: 5 s. The parameters for sine wave illumination were as follows: amplitude: typically 1 mW; frequency: 1, 4, 8, 12, and 20 Hz; number of trials: 20; minimum number of cycles at each trial: 10; minimum trial duration: 5 s. Illumination was conducted through a 450 nm laser diode (PL450B, Osram) driven via a current controller (LDC205C; Thorlabs) and a data acquisition board (USB-6212, National Instruments). DiD solution (2% in ethanol, Thermo Fisher Scientific Cat# D307, Waltham, MA, USA) was applied to the non-recording side of the shank of optoprobes for post hoc location of the recording sites.

#### Seizure interventions via MS stimulation

##### Electrical and optogenetic stimulation

During the chronic implantation surgery of the recording electrodes, either a bipolar stimulating electrode (same construct as the kindling electrode) or an optical cannula targeting the MS was implanted, too. The coordinates used were as follows: AP: 0.5 mm anterior from the bregma; ML: penetrated from 1.0 mm leftward from the midline at a 9° leftward tilted angle from the parasagittal plane; Distance: 6.0 mm from the pia. The stimulating electrode or the optical cannula was fixed with dental cement. MS stimuli for seizure interventions were commanded by TTL signals either before (pre-conditioning) or after (closed-loop or open-loop) seizure induction by E-stim of the HPC commissure. For E-stim of the MS, each TTL signal triggered a bipolar, 0.5 ms positive, 0.5 ms negative rectangular pulse at ±400 μA intensity. For optogenetic stimulation, each TTL signal triggered 30 ms-long square wave illumination at maximum intensity through each optical cannula (typically 30–40 mW).

##### Open-loop septal stimulation experiments

Either electrical or optogenetic MS stimuli were delivered for two minutes at a fixed frequency (1, 8, or 20 Hz) following each seizure induction. These stimulus frequencies were chosen because low- and theta-frequency stimulation of the limbic system during interictal periods alleviate temporal lobe epilepsy (42, 69) and 20 Hz was the maximum frequency MS stimulation at which frequency did not cause seizures in non-kindled rats in pilot experiments. Intensity of E-stim for seizure induction was determined either as the minimum intensity by which HPC after-discharges were reliably (>90%) induced (supra-threshold stimulation: ±10–100 μA typically) or the maximum intensity by which HPC after-discharges were seldom (<10%) induced (sub-threshold stimulation: ±5–50 μA typically). MS stimulated and non-stimulated trials were always alternated, with the same seizure induction intensity in a pseudo random manner.

##### Closed-loop septal stimulation experiments

HPC electrographic seizure waves were detected on-line using a programmable digital signal processor unit (RX-8; Tucker-Davis Technologies, Alachua, FL, USA) to trigger either electrical or optogenetic stimulation of the MS for on-demand real-time seizure interventions. Pre-amplified and multiplexed LFP signals were fed to the processor unit in parallel to the recording unit sampled at 500 Hz, and the signals were demultiplexed and analysed on-line to detect each population spike in HPC LFPs using a custom-made seizure detection algorithm based on a previously established one (23). Briefly, LFP signals were demultiplexed at 500 Hz per channel and a signal from a pre-selected HPC channel was band-pass filtered with a 4^th^ order Butterworth filter to 10–130 Hz. The detection was triggered if the filtered signal consequently exceeded a threshold within a time window (typically 30–300 ms). Both threshold for amplitude and a duration of the time-window were fine-turned for each animal. A delay (0, 20, 40, or 60 ms) was introduced between detection and stimulation to target a specific phase of seizure waves (Figure 1). Intensity of E-stim for seizure induction was determined as a minimum intensity by which HPC after-discharges were reliably (>90%) induced (supra-threshold stimulation: ±10–100 μA typically). MS stimulated and non-stimulated trials were always paired with the same seizure induction intensity in a pseud random manner. Trials in which no after-discharges were observed were discarded.

##### Pre-conditioning

Either electrical or optogenetic MS stimuli were delivered for 5 or 60 seconds at 20 Hz before seizure induction. Intensity of seizure induction were varied trial-by-trial to determine thresholds for HPC and Ctx electrographic seizures: regenerative after-discharges in the HPC and large amplitude oscillations in cortical LFPs associated with Racine’s scale 4 or 5. The stimulus intensity was typically started from ±10 μA and incremented at ±10 μA steps. MS stimulated and non-stimulated trials were always paired with a pseudo random sequence of stimulus intensity.

#### Immunohistochemistry

##### Colchicine injection

To enhance immunohistochemical reaction of cellular markers (glutamate and GAD), 150 μg colchicine dissolved in saline (C9754; Sigma-Aldrich) was pressure-injected into the lateral ventricle 48 h before perfusion (38). The stereotaxic coordinate of the injection site was as follows: AP, 0.8 mm posterior from the bregma; ML, 1.5 mm rightward from the midline; DV, 4.4 mm from the pia.

##### Perfusion and sectioning

The animals were deeply anesthetized with 1.5 g/kg urethane (i.p.) and transcardially perfused with physiological saline followed by 4% paraformaldehyde (PFA) and 0.2% picric acid (PA) in 0.1M phosphate buffer (PB) (pH 7.2–7.3). After removal, brains were postfixed in the same fixative orvernight, embedded into 4% agarose, and sectioned in 40–50 μm thick slices using a vibrating blade microtome (VT1000S, Leica, Buffalo Grove, IL, USA). Sections were collected in 0.1 M phosphate buffered saline (PBS), and then subjected to staining described below. All staining procedures were performed at room temperature.

##### NeuN immunohistochemistry

All incubations were followed by washing with 0.3% Triton-X in 0.1 M PBS (PBS-X). Sections were incubated successively with 10% normal goat serum (NGS) in PBS-X for 30 min, 1:1000 diluted mouse anti-NeuN monoclonal antibody (Millipore Cat# MAB377, RRID:AB_2298772) in PBS-X containing 1% NDS and 0.02% sodium azide (PBS-XG) overnight, and 2 μg/ml Alexa Fluor 488-conjugated goat anti-mouse IgG for two hours (Thermo Fisher Scientific Cat# A-11029, RRID:AB_2534088). Sections were then counter-stained with fluoro-Nissl solution (Thermo Fisher Scientific Cat# N-21479). Sections were finally mounted on gelatine-coated glass slides, cover-slipped with 50% glycerol and 2.5% triethylene diamine in PBS.

##### GAD-67/65-immunohistochemistry

Somatic GAD expression levels were enhanced by colchicine as described above. Sections were incubated successively with 10% NGS in PBS-X for 30 min, 1 μg/ml rabbit anti-GAD67/65 polyclonal antibody (Frontier Institute Cat# GAD-Rb, RRID:AB_2571698) in PBS-XG overnight, and 2 μg/ml Alexa Fluor 633-conjugated goat anti-Rabbit IgG for two hours (Thermo Fisher Scientific Cat# A-21071, RRID:AB_2535732).

##### Glutamate immunohistochemistry

Somatic glutamate expression levels were enhanced by colchicine as described above. The animals were perfused as described above but 0.5% glutaraldehyde was added to the fixative solution. Sections were incubated successively with 10% normal donkey serum (NDS) in PBS-X for 30 min, 1:4000 diluted mouse anti-glutamate monoclonal antibody (ImmunoStar Cat# 22523, RRID:AB_572244) in PBS-X containing 1% NDS and 0.02% sodium azide (PBS-XD) overnight, and 2 μg/ml Alexa Fluor 647-conjugated donkey anti-mouse IgG for two hours (Thermo Fisher Scientific Cat# A-31571, RRID:AB_162542).

##### ChAT-immunohistochemistry

Sections were incubated successively with 10% NDS in PBS-X for 30 min, 1:500 diluted goat anti-ChAT polyclonal antibody (Millipore Cat# AB144P, RRID:AB_2079751) in PBS-XD overnight, and 2 μg/ml Alexa Fluor 633-conjugated donkey anti-goat IgG in PBS-XD for two hours (Thermo Fisher Scientific Cat# A-21082, RRID:AB_2535739).

##### cFos-immunohistochemistry

Just before perfusion, the MS with ChR2 transduction were laser-illuminated with 20 Hz sinusoidal waves (20 mW) for two hours in freely-moving states as described above. Sections were incubated successively with 10% NGS in PBS-X for 30 min, 1:4000 diluted rabbit anti-cFos polyclonal antibody (Millipore Cat# ABE457, RRID:AB_2631318) in PBS-XG overnight, and 4 μg/ml Alexa Fluor 633-conjugated goat anti-rabbit IgG in PBS-XG for two hours (Thermo Fisher Scientific).

##### Concurrent GAD, Glutamate, and ChAT immunohistochemistry

Somatic expression levels were enhanced by colchicine as described above. Sections were incubated successively with 10% NDS in PBS-X for 30 min, a mixture of primary antibodies in PBS-XD overnight (1 μg/ml rabbit anti-GAD67/65, Frontier; 1:4000 mouse anti-glutamate, ImmunoStar; 1:500 goat anti-ChAT, Millipore), and a mixture of secondary antibodies in PBX-XD for two hours (donkey anti-goat, rabbit, and mouse IgGs conjugated with Alexa Fluor 488, 555, and 647, respectively; 2 μg/ml each; Thermo Fisher Scientific Cat# A-11055, RRID:AB_2534102; Cat# A-31572, RRID:AB_162543, and Cat# A-31571, RRID:AB_162542).

##### GAD and PV double immunohistochemistry

Somatic expression levels were enhanced by colchicine as described above. Sections were incubated successively with 10% NDS in PBS-X for 30 min, a mixture of primary antibodies in PBS-XD overnight (1 μg/ml rabbit anti-GAD67/65, Frontier and 1:3000 mouse anti-PV, Swant Cat# 235, RRID: AB_10000343), and a mixture of secondary antibodies in PBS-XD for two hours (Alexa Fluor 555-conjugated donkey anti-rabbit IgG and Alexa 647 conjugated donkey anti-mouse IgG; 2 μg/ml, Thermo Fisher Scientific).

#### Confocal microscopy

Stained sections were examined using a Zeiss LSM880 scanning confocal microscope (Carl Zeiss, Oberkochen, Germany). Images of 1–2 μm-optical thickness were acquired using a Plan-Apochromat 40×/1.4 Oil DIC M27 or an alpha Plan-Apochromat 63×/1.46 Oil Korr M27 objective lens, 4.12 μs pixel time, and 16 times frame average at 512 × 512 resolution. The illumination power ranged 0.4–0.7 mW, which did not induce obvious fading.

### QUANTIFICATION AND STATISTICAL ANALYSIS

All data analyses were performed in MATLAB (RRID:SCR_001622; Mathworks, Natick, MA, USA), unless otherwise noted.

#### Duration of electrographic seizures

Wide-band 20 k sample/s signals of open-loop intervention experiments were first down-sampled to 1250 sample/s and filtered with a 3^rd^ order zero phase lag Butterworth filter between 1 and 625 Hz to prepare LFP signals. Signals of closed-loop intervention experiments sampled at 500 Hz were filtered between 1 and 250 Hz for LFP signals. Peri-stimulus LFPs (30 s baseline and 180 s test epochs) were then extracted using timestamps recorded in a digital channel. The peri-stimulus LFP signals were further band-pass filtered to 10–80 Hz to prepare narrow-band LFPs. The narrow-band LFPs were then smoothed using a three-second-long moving average filter. Durations of the HPC and Ctx electrographic seizures were automatically detected and defined as duration in test epochs when all amplitude of the smoothed LFPs in each brain region exceeded three times the root-mean-square levels of the corresponding baseline epochs. Duration of electrographic seizures with MS electrical interventions were refined by the consensus estimate of manual inspections by two experienced researchers using Neuroscope software (RRID:SCR_002455) (70) because the automated-detection algorithm sometimes misestimated due to the electrical artifacts. Coefficient of variation between manual and automated detections examined with data of optogenetic interventions was less than five percent on both researchers.

#### Modulation index of HPC electrographic seizures

Modulation index (MI) of duration of HPC electrographic seizures by MS stimulation was defined as (HPC seizure duration with MS stimulation − HPC seizure duration without MS stimulation) / (HPC seizure duration with MS stimulation + HPC seizure duration without MS stimulation). Distributions of MIs were fitted with a Gaussian mixture model for clustering of possible sub-populations with anti-seizure effects (‘Success’ and ‘Non-success’ trials) or pro-seizure effects (‘Induction’ or ‘Non-induction’ trials). Skewness (asymmetry) of MI distributions was evaluated by comparing absolute value distributions of positive and negative MIs.

#### Assessment of severity of motor seizures

Severity of motor seizures was video monitored and Hilbert transform and significant distribution of evaluated according to the Racine’s scale on each each unit on the phase was tested using the trial (47).

#### Spike sorting and unit classification

Neuronal spikes were detected from the digitally high-pass filtered raw signal (0.5–5 kHz) by a threshold crossing-based algorithm and detected spikes were automatically sorted using a masked EM algorithm for Gaussians mixtures (71) implemented in Klusta software, an open source automatic spike sorting package (https://github.com/kwikteam/klusta) (Dataset figure 5). This automatic clustering process was followed by manual refinement of the clusters using KlustaViewa software to obtain well-isolated single units (72) (https://github.com/klusta-team/klustaviewa/).

Multiunit and noise clusters were discarded during the manual process. Quality of cluster isolation was estimated by calculating the isolation distance and interspike interval index for each cluster as previously described (66); poor quality clusters were discarded.

#### Detection of optogenetically modulated units

For detection of modulated units by 10 ms pulse illumination, peri-stimulus time histogram (PSTH) was prepared and cross-correlation (CCG) analysis has been applied between stimulus and spike timestamps (Dataset figure 5). Significant modulation in firing rate of each unit was identified using a shuffling method. For each stimulus and unit pair, surrogate data set of timestamps were constructed by random permutation of labels: either stimulus or unit 200 times. Point-wise 95% acceptance bands were calculated from the surrogate data set CCGs for each 1 ms bin and multiple comparison error was corrected by introducing ‘global significance bands’ (73). The 10 ms illumination was considered to have excitatory or inhibitory effects on the referred unit if any of its CCG bins reached above or below these global bands within the considered time window.

For detection of modulated units by sinusoidal illumination, instantaneous phase values of stimulus wave were calculated using Hilbert transform and significant distribution of each unit on the phase was tested using the Rayleigh-test in CircStat MATLAB Toolbox with a 0.05 alpha level (74).

#### Detection of modulated units by septal LFPs

LFP signals were calculated on each recording site in the MS by 1250 Hz down-sampling and <300 Hz low-pass filtering and the LFP signals were averaged (Dataset figure 5). The averaged LFP signal was then band-pass filtered into five frequency bands (delta, 1–4 Hz; theta, 4–12 Hz; beta, 12–30 Hz; low-gamma, 30–45 Hz; mid-gamma, 45–80 Hz; high-gamma, 80–150 Hz) with an 3^rd^ order zero phase-lag Butterworth filter, and instantaneous phase values were calculated using Hilbert-transform. Significant modulations of each unit by these LFP frequency bands were tested using the Rayleigh-test.

#### Image analysis

Acquired raw image files were opened and ChR2-mCherry and immunohistochemical signals were automatically and linearly enhanced with a Carl Zeiss software (ZEN Digital Imaging for Light Microscopy, RRID:SCR_013672), and each image was exported as an eight-bit tiff file. Colocalizations between ChR2-mCherry signals and immunohistochemical reactions of cell-type markers were manually inspected with Cell Counter plugin of ImageJ software (RRID:SCR_003070, NIH, Bethesda, MD, USA).

#### Statistical analysis

Values are given as mean ± standard deviation (s.d.) unless otherwise noted. MATLAB with Statistics and Machine Learning Toolbox was used for statistical tests. Two-tailed paired *t*-test was employed to compare means of durations of electrographic seizures and threshold intensity of seizure induction w/wo septal pre-conditioning. Wilcoxon singed rank test was employed to compare the Racine’s scales between trials w/wo MS stimulation. χ^2^ test was employed to examine biased fractions of MI-based clustering (Seizure induction trials or Success trials) over a MS stimulus condition (frequency or delay). Two-way repeated measure ANOVA was employed to test interaction of stimulus frequency or stimulus delay and main effect of MS stimulation. Three-way repeated measure ANOVA was employed to examine consistent interactions between MS stimulation and clustering based on MI on seizure parameters (HPC seizure duration, Ctx seizure duration, and Racine’s scale). Spearman’s rank correlation test was employed to examine correlations between MI of HPC seizure duration, changes of Ctx seizure duration, and changes in Racine’s scale by MS stimulation. Two-sample Kolmogorov-Smirnov test was employed to compare absolute value distributions of positive and negative MIs and distributions of HPC seizure durations w/wo MS stimulation to evaluate skewness. The significance level was set at *P* < 0.05. One, two, and three asterisks on the figures denote significance levels <0.05, <0.01, and <0.001, respectively.

## DATA AND SOFTWARE AVAILABILITY

All of the raw data and codes used to generate the graphs shown in the figures and additional dataset figures are available at http://www.data.mendeley.com (ID 10.17632/k9hwm7p33x.1).

## References

1. Kwan P, Schachter SC, Brodie MJ. Drug-Resistant Epilepsy. New Engl J Med. 2011;365(10):919–26.

2. Chen Z, Brodie MJ, Liew D, Kwan P. Treatment Outcomes in Patients With Newly Diagnosed Epilepsy Treated With Established and New Antiepileptic Drugs: A 30-Year Longitudinal Cohort Study. JAMA Neurol. 2018;75(3):279–86.

3. Bone B, Fogarasi A, Schulz R, Gyimesi C, Kalmar Z, Kovacs N, et al. Secondarily generalized seizures in temporal lobe epilepsy. Epilepsia. 2012;53(5):817–24.

4. Massey CA, Sowers LP, Dlouhy BJ, Richerson GB. Mechanisms of sudden unexpected death in epilepsy: the pathway to prevention. Nat Rev Neurol. 2014;10(5):271–82.

5. Holmes MD, Miles AN, Dodrill CB, Ojemann GA, Wilensky AJ. Identifying Potential Surgical Candidates in Patients with Evidence of Bitemporal Epilepsy. Epilepsia. 2003;44(8):1075–9.

6. Berg AT, Vickrey BG, Langfitt JT, Sperling MR, Walczak TS, Shinnar S, et al. The Multicenter Study of Epilepsy Surgery: Recruitment and Selection for Surgery. Epilepsia. 2003;44(11):1425–33.

7. Selwa LM, Schmidt SL, Malow BA, Beydoun A. Long‐term Outcome of Nonsurgical Candidates with Medically Refractory Localization‐related Epilepsy. Epilepsia. 2003;44(12):1568–72.

8. Li MCH, Cook MJ. Deep brain stimulation for drug‐resistant epilepsy. Epilepsia. 2018;59(2):273–90.

9. Han C-L, Hu W, Stead M, Zhang T, Zhang J-G, Worrell GA, et al. Electrical stimulation of hippocampus for the treatment of refractory temporal lobe epilepsy. Brain Res Bull. 2014;109:13–21.

10. Cukiert A, Lehtimäki K. Deep brain stimulation targeting in refractory epilepsy. Epilepsia. 2017;58(S1):80–4.

11. Paz JT, Huguenard JR. Microcircuits and their interactions in epilepsy: is the focus out of focus? Nat Neurosci. 2015;18(3):351–9.

12. Fisher R, Salanova V, Witt T, Worth R, Henry T, Gross R, et al. Electrical stimulation of the anterior nucleus of thalamus for treatment of refractory epilepsy. Epilepsia. 2010;51(5):899–908.

13. Fountas KN, Kapsalaki E, Hadjigeorgiou G. Cerebellar stimulation in the management of medically intractable epilepsy: a systematic and critical review. Neurosurg Focus. 2010;29(2):E8.

14. Dutar P, Bassant MH, Senut MC, Lamour Y. The septohippocampal pathway: structure and function of a central cholinergic system. Physiol Rev. 1995;75(2):393–427.

15. Kang D, Ding M, Topchiy I, Kocsis B. Reciprocal Interactions between Medial Septum and Hippocampus in Theta Generation: Granger Causality Decomposition of Mixed Spike-Field Recordings. Front Neuroanat. 2017;11:120.

16. Buzsáki G. Theta oscillations in the hippocampus. Neuron. 2002;33(3):325–40.

17. Vandecasteele M, Varga V, Berényi A, Papp E, Barthó P, Venance L, et al. Optogenetic activation of septal cholinergic neurons suppresses sharp wave ripples and enhances theta oscillations in the hippocampus. Proc Natl Acad Sci USA. 2014;111(37):13535–40.

18. Malow BA, Carney PR, Kushwaha R, Bowes RJ. Hippocampal sleep spindles revisited: physiologic or epileptic activity? Clin Neurophysiol. 1999;110(4):687–93.

19. Ewell LA, Liang L, Armstrong C, Soltesz I, Leutgeb S, Leutgeb JK. Brain State Is a Major Factor in Preseizure Hippocampal Network Activity and Influences Success of Seizure Intervention. J Neurosci. 2015;35(47):15635–48.

20. Moxon KA, Shahlaie K, Girgis F, Saez I, Kennedy J, Gurkoff GG. From adagio to allegretto: The changing tempo of theta frequencies in epilepsy and its relation to interneuron function. Neurobiol Dis. 2019;129(Cell Tissue Res. 258 1989):169–81.

21. Morrell MJ. Responsive cortical stimulation for the treatment of medically intractable partial epilepsy. Neurology. 2011;77(13):1295–304.

22. Meiron O, Gale R, Namestnic J, Bennet-Back O, Gebodh N, Esmaeilpour Z, et al. Antiepileptic Effects of a Novel Non-invasive Neuromodulation Treatment in a Subject With Early-Onset Epileptic Encephalopathy: Case Report With 20 Sessions of HD-tDCS Intervention. Front Neurosci. 2019;13:547.

23. Kozák G, Berényi A. Sustained efficacy of closed loop electrical stimulation for long-term treatment of absence epilepsy in rats. Sci Rep. 2017;7(1):6300.

24. Berényi A, Belluscio M, Mao D, Buzsáki G. Closed-Loop Control of Epilepsy by Transcranial Electrical Stimulation. Science. 2012;337(6095):735–7.

25. Dejean C, Courtin J, Karalis N, Chaudun F, Wurtz H, Bienvenu TCM, et al. Prefrontal neuronal assemblies temporally control fear behaviour. Nature. 2016;535(7612):420–4.

26. Tsanov M. Differential and complementary roles of medial and lateral septum in the orchestration of limbic oscillations and signal integration. Eur J Neurosci. 2018;48(8):2783–94.

27. Sinel’nikova VV, Popova YI, Kichigina VF. Correlational Relationships Between the Hippocampus and Medial Septal Area and Their Changes During Epileptogenesis. Neurosci Behav Physiol. 2009;39(7):619–23.

28. Fuhrmann F, Justus D, Sosulina L, Kaneko H, Beutel T, Friedrichs D, et al. Locomotion, Theta Oscillations, and the Speed-Correlated Firing of Hippocampal Neurons Are Controlled by a Medial Septal Glutamatergic Circuit. Neuron. 2015;86(5):1253–64.

29. Robinson J, Manseau F, Ducharme G, Amilhon B, Vigneault E, El Mestikawy S, et al. Optogenetic Activation of Septal Glutamatergic Neurons Drive Hippocampal Theta Rhythms. J Neurosci. 2016;36(10):3016–23.

30. Zutshi I, Brandon MP, Fu ML, Donegan ML, Leutgeb JK, Leutgeb S. Hippocampal Neural Circuits Respond to Optogenetic Pacing of Theta Frequencies by Generating Accelerated Oscillation Frequencies. Curr Biol. 2018;28(8):1179–88.

31. McIntyre DC, Gilby KL. Mapping seizure pathways in the temporal lobe. Epilepsia. 2008;49 Suppl 3:23–30.

32. Joshi A, Salib M, Viney T, Dupret D, Somogyi P. Behavior-Dependent Activity and Synaptic Organization of Septo-hippocampal GABAergic Neurons Selectively Targeting the Hippocampal CA3 Area. Neuron. 2017;96(6):1342–57.

33. Colom LV, Castaneda MT, Reyna T, Hernandez S, Garrido-Sanabria E. Characterization of Medial Septal Glutamatergic Neurons and Their Projection to the Hippocampus. Synapse. 2005;58(3):151–64.

34. Desikan S, Koser DE, Neitz A, Monyer H. Target selectivity of septal cholinergic neurons in the medial and lateral entorhinal cortex. Proc Natl Acad Sci USA. 2018;115(11):E2644–E52.

35. Chamberland S, Salesse C, Topolnik D, Topolnik L. Synapse-specific inhibitory control of hippocampal feedback inhibitory circuit. Front Cell Neurosci. 2010;4:130.

36. Gelinas JN, Khodagholy D, Thesen T, Devinsky O, Buzsáki G. Interictal epileptiform discharges induce hippocampal-cortical coupling in temporal lobe epilepsy. Nat Med. 2016;22(6):641–8.

37. Monai H, Ohkura M, Tanaka M, Oe Y, Konno A, Hirai H, et al. Calcium imaging reveals glial involvement in transcranial direct current stimulation-induced plasticity in mouse brain. Nat Commun. 2016;7(1):11100.

38. Gritti I, Manns ID, Mainville L, Jones BE. Parvalbumin, calbindin, or calretinin in cortically projecting and GABAergic, cholinergic, or glutamatergic basal forebrain neurons of the rat. J Comp Neurol. 2003;458(1):11–31.

39. Beekwilder JP, Beems T. Overview of the Clinical Applications of Vagus Nerve Stimulation. J Clin Neurophysiol. 2010;27(2):130–8.

40. Müller C, Remy S. Septo–hippocampal interaction. Cell Tissue Res. 2017;373(3):565–75.

41. Izadi A, Pevzner A, Lee DJ, Ekstrom AD, Shahlaie K, Gurkoff GG. Medial Septal Stimulation Increases Seizure Threshold and Improves Cognition in Epileptic Rats. Brain Stimul. 2019;12(3):735–42.

42. Miller JW, Turner GM, Gray BC. Anticonvulsant effects of the experimental induction of hippocampal theta activity. Epilepsy Res. 1994;18(3):195–204.

43. Colom LV, García-Hernández A, Castañeda MT, Perez-Cordova MG, Garrido-Sanabria ER. Septo-hippocampal networks in chronically epileptic rats: potential antiepileptic effects of theta rhythm generation. J Neurophysiol. 2006;95(6):3645–53.

44. Soares JI, Costa C, Ferreira MH, Andrade PA, Maia GH, Lukoyanov NV. Partial depletion of septohippocampal cholinergic cells reduces seizure susceptibility, but does not mitigate hippocampal neurodegeneration in the kainate model of epilepsy. Brain Res. 2019;1717:235–46.

45. Paxinos G, Watson C. The Rat Brain in Stereotaxic Coordinates. 6th ed. New York: Academic Press; 2006.

46. Tsanov M. Speed and Oscillations: Medial Septum Integration of Attention and Navigation. Front Syst Neurosci. 2017;11:67.

47. Racine RJ. Modification of seizure activity by electrical stimulation. II. Motor seizure. Electroencephalogr Clin Neurophysiol. 1972;32(3):281–94.

48. Beudel M, Brown P. Adaptive deep brain stimulation in Parkinson's disease. Parkinsonism Relat Disord. 2016;22:S123–S6.

49. Jirsa VK, Stacey WC, Quilichini PP, Ivanov AI, Bernard C. On the nature of seizure dynamics. Brain. 2014;137(8):2210–30.

50. Kalitzin S, Petkov G, Suffczynski P, Grigorovsky V, Bardakjian BL, Lopes da Silva F, et al. Epilepsy as a manifestation of a multistate network of oscillatory systems. Neurobiol Dis. 2019;130:104488.

51. Nordholm S, Schepker H, Tran LTT, Doclo S. Stability-controlled hybrid adaptive feedback cancellation scheme for hearing aids. J Acoust Soc Am. 2018;143(1):150–66.

52. Unal G, Joshi A, Viney TJ, Kis V, Somogyi P. Synaptic Targets of Medial Septal Projections in the Hippocampus and Extrahippocampal Cortices of the Mouse. J Neurosci. 2015;35(48):15812–26.

53. Hájos N, Paulsen O. Network mechanisms of gamma oscillations in the CA3 region of the hippocampus. Neural Netw. 2009;22(8):1113–9.

54. Cossart R. Operational hub cells: a morpho-physiologically diverse class of GABAergic neurons united by a common function. Curr Opin Neurobiol. 2014;26:51–6.

55. Owen SF, Liu MH, Kreitzer AC. Thermal constraints on in vivo optogenetic manipulations. Nat Neurosci. 2019;22(7):1061–5.

56. Freund TF, Antal M. GABA-containing neurons in the septum control inhibitory interneurons in the hippocampus. Nature. 1988;336(6195):170–3.

57. Gulyás AI, Freund TT. Generation of physiological and pathological high frequency oscillations: the role of perisomatic inhibition in sharp-wave ripple and interictal spike generation. Curr Opin Neurobiol. 2015;31:26–32.

58. Haam J, Zhou J, Cui G, Yakel JL. Septal cholinergic neurons gate hippocampal output to entorhinal cortex via oriens lacunosum moleculare interneurons. Proc Natl Acad Sci USA. 2018;115(8):E1886–E95.

59. Fisher RS. Stimulation of the medial septum should benefit patients with temporal lobe epilepsy. Med Hypotheses. 2015;84(6):543–50.

60. Izadi A, Ondek K, Schedlbauer A, Keselman I, Shahlaie K, Gurkoff G. Clinically indicated electrical stimulation strategies to treat patients with medically refractory epilepsy. Epilepsia Open. 2018;3(Suppl Suppl 2):198–209.

61. Neumann W-J, Turner RS, Blankertz B, Mitchell T, Kühn AA, Richardson MR. Toward Electrophysiology-Based Intelligent Adaptive Deep Brain Stimulation for Movement Disorders. Neurotherapeutics. 2019;16(1):105–18.

62. Takeuchi Y, Berényi A. Oscillotherapeutics – Time-targeted interventions in epilepsy and beyond. Neurosci Res. 2020:in press (doi: 10.1016/j.neures.2020.01.002).

63. Witten IB, Steinberg EE, Lee SY, Davidson TJ, Zalocusky KA, Brodsky M, et al. Recombinase-driver rat lines: tools, techniques, and optogenetic application to dopamine-mediated reinforcement. Neuron. 2011;72(5):721–33.

64. Iwasato T, Nomura R, Ando R, Ikeda T, Tanaka M, Itohara S. Dorsal telencephalon-specific expression of Cre recombinase in PAC transgenic mice. Genesis. 2004;38(3):130–8.

65. Kozák G, Földi T, Berényi A. Chronic Transcranial Electrical Stimulation and Intracortical Recording in Rats. J Vis Exp. 2018(135):e56669.

66. Berényi A, Somogyvári Z, Nagy AJ, Roux L, Long JD, Fujisawa S, et al. Large-scale, high-density (up to 512 channels) recording of local circuits in behaving animals. J Neurophysiol. 2014;111(5):1132–49.

67. Lerner-Natoli M, Rondouin G, Baldy-Moulinier M. Evolution of Wet Dog Shakes during Kindling in Rats: Comparison between Hippocampal and Amygdala Kindling. Exp Neurol. 1984;83(1):1–12.

68. Royer S, Zemelman BV, Barbic M, Losonczy A, Buzsáki G, Magee JC. Multi-array silicon probes with integrated optical fibers: light-assisted perturbation and recording of local neural circuits in the behaving animal. Eur J Neurosci. 2010;31(12):2279–91.

69. Wang Y, Liang J, Xu C, Wang Y, Kuang Y, Xu Z, et al. Low-frequency stimulation in anterior nucleus of thalamus alleviates kainate-induced chronic epilepsy and modulates the hippocampal EEG rhythm. Exp Neurol. 2016;276:22–30.

70. Hazan L, Zugaro M, Buzsáki G. Klusters, NeuroScope, NDManager: A free software suite for neurophysiological data processing and visualization. J Neurosci Methods. 2006;155(2):207–16.

71. Kadir SN, Goodman DF, Harris KD. High-Dimensional Cluster Analysis with the Masked EM Algorithm. Neural comput. 2014;26(11):2379–94.

72. Rossant C, Kadir SN, Goodman DF, Schulman J, Hunter ML, Saleem AB, et al. Spike sorting for large, dense electrode arrays. Nat Neurosci. 2016;19(4):634–41.

73. Fujisawa S, Amarasingham A, Harrison MT, Buzsáki G. Behavior-dependent short-term assembly dynamics in the medial prefrontal cortex. Nat Neurosci. 2008;11(7):823–33.

74. Berens P. CircStat: A MATLABToolbox for Circular Statistics. J Statisticcal Software. 2009;31(10):10

