## Supplementary material for "Closed-loop stimulation of the medial septum terminates epileptic seizures": Figure S1-7, Table S2

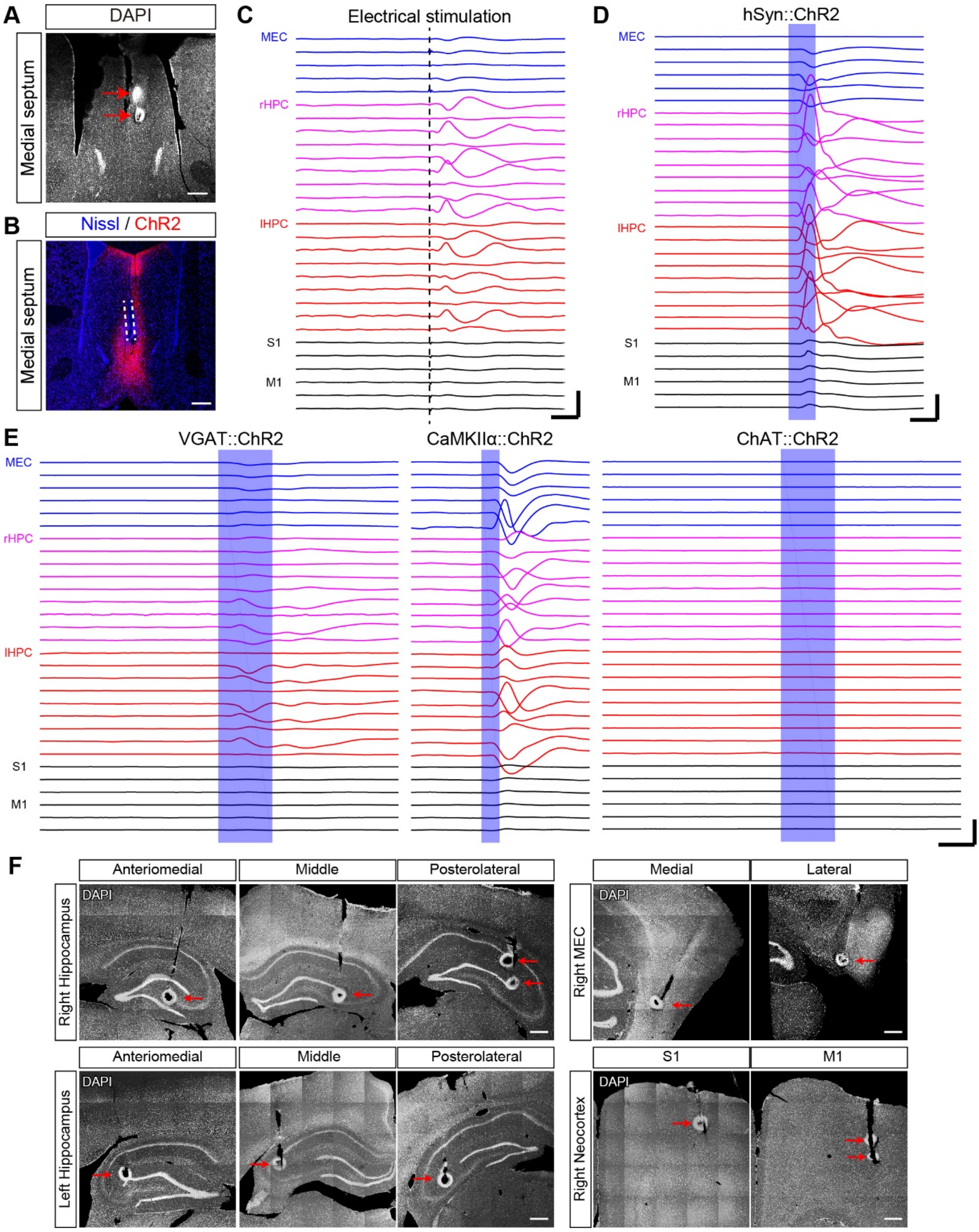


**Figure S1. Electrical and optogenetic stimulation of the MS evoked LFP deflections in the HPC of awake freely-moving rats, Related to Figure 1**

(**A**, **B**) Stimulation sites in the MS. Red arrows indicate electrically lesioned sites via a bipolar stimulus electrode (A). Dashed lines indicate the track of optical cannula insertion in the MS with AAV-mediated ChR2-mCherry transduction (B). Scale bars: 0.5 mm. (**C**, **D**) Averaged LFPs evoked by electrical (C) and non-specific optogenetical (D) stimulation of the MS. Dashed vertical line and a blue patch represent stimulation timing for (C) and (D), respectively. One hundred to one thousand trials were averaged for each trace. Scale bars: 2 mV, 20 ms. (**E**) The same conventions as (C) and (D) for selective GABAergic (left), glutamatergic (middle), and cholinergic (right) optical stimulation. (**F**) Post-mortem location of recording sites. One (the deepest) or two (deepest and middle) recording sites on each shank were lesioned by direct anodal currents via tips. Each arrow indicates each recording site. Scale bars: 0.5 mm.

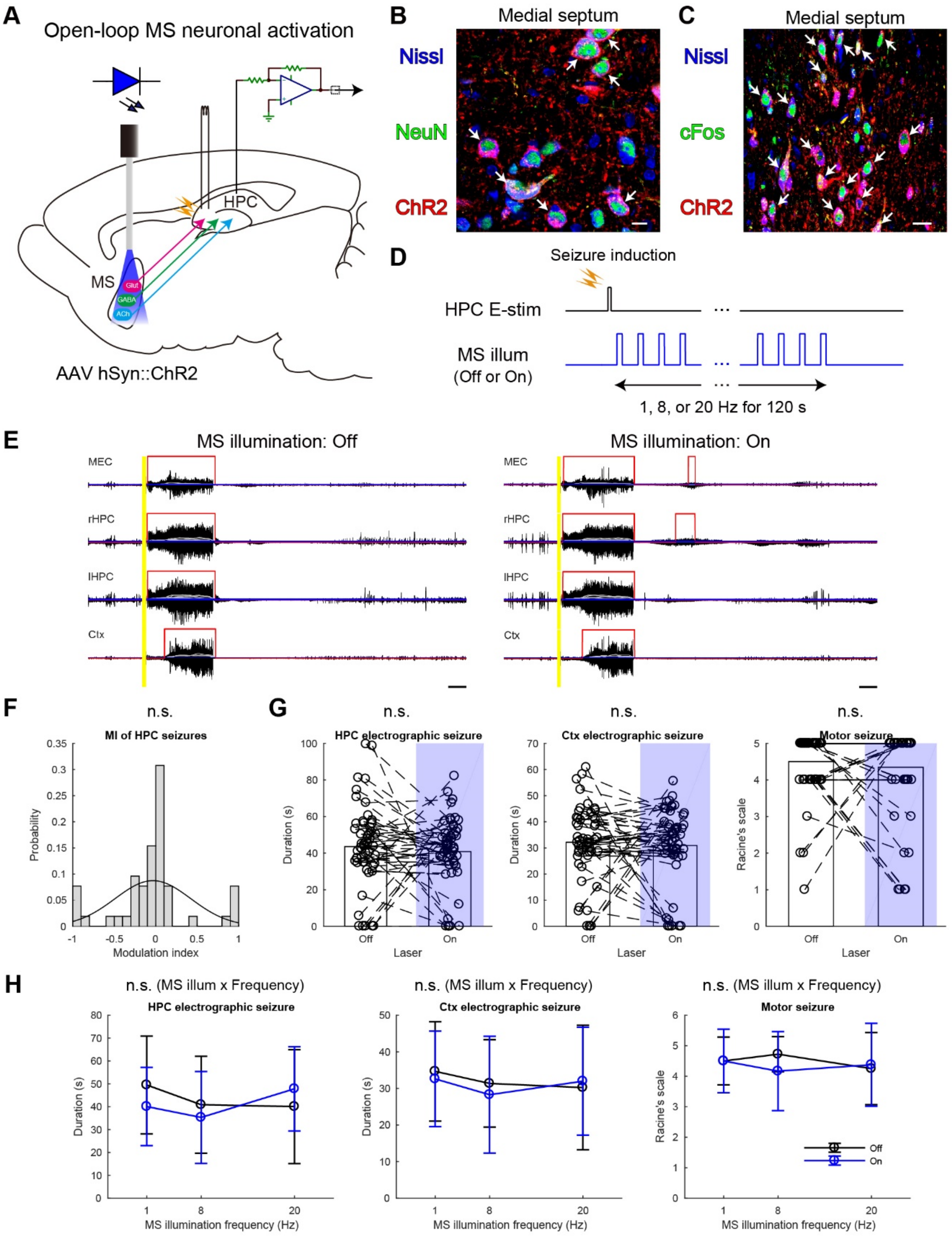
A
Open-loop MS neuronal activation
HPC
MS
Glut
GABA
ACh
AAV hSyn::ChR2
B
Medial septum
Nissl
NeuN
ChR2
C
Medial septum
Nissl
cFos
ChR2
D
Seizure induction
HPC E-stim
MS illum
(Off or On)
1, 8, or 20 Hz for 120 s
E
MS illumination: Off
MS illumination: On
MEC
rHPC
lHPC
Ctx
F
n.s.
MI of HPC seizures
Probability
Modulation index
G
n.s.
HPC electrographic seizure
Ctx electrographic seizure
Motor seizure
Duration (s)
Racine's scale
Off
On
Laser
H
n.s. (MS illum x Frequency)
HPC electrographic seizure
Ctx electrographic seizure
Motor seizure
Duration (s)
Racine's scale
MS illumination frequency (Hz)
Off
On

**Figure S2. Open-loop optogenetic activation of MS neurons does not alleviate seizures of HPC-origin, Related to Figure 2**

(**A**) Schema of the experiment. A stimulus electrode and an optical cannula were inserted into the HPC and the MS for seizure induction and intervention, respectively. (**B**) Expression of ChR2 in the MS. ChR2-mCherry signals were colocalized with NeuN immuno-positive neurons. Each arrow indicates the colocalization. Scale bar: 10 μm. (**C**) cFos immunohistochemical reactions were colocalized with ChR2-mCherry expressing MS neurons after illumination. Each arrow indicates the colocalization. Scale bar: 20 μm. (**D**) Concept of open-loop responsive optogenetic seizure intervention. (**E**) Representative seizure waves w/wo open-loop MS illumination. The same conventions were used as in Figure 2C. (**F**) Distribution of MIs with MS illumination. Curve represents a fitted Gaussian model. (**G**) Duration of HPC and Ctx seizures, and motor seizures. Data of different illumination frequencies were pooled. Bars represent means, while circles represent each individual trial pairs. (**H**) Illumination frequency resolved representation of the data shown in (G). Values are represented as means ± s.d. n = 104 trials from four rats. Statistical significance was tested by two-sample Kolmogorov-Smirnov for asymmetry of MI distributions (D, F), paired *t*-test for seizure durations and Wilcoxon singed rank test for the Racine's scale (G) and tested by two-way repeated ANOVA for (H). Results of statistical tests are extensively reported in Table S1. n.s., not significant.

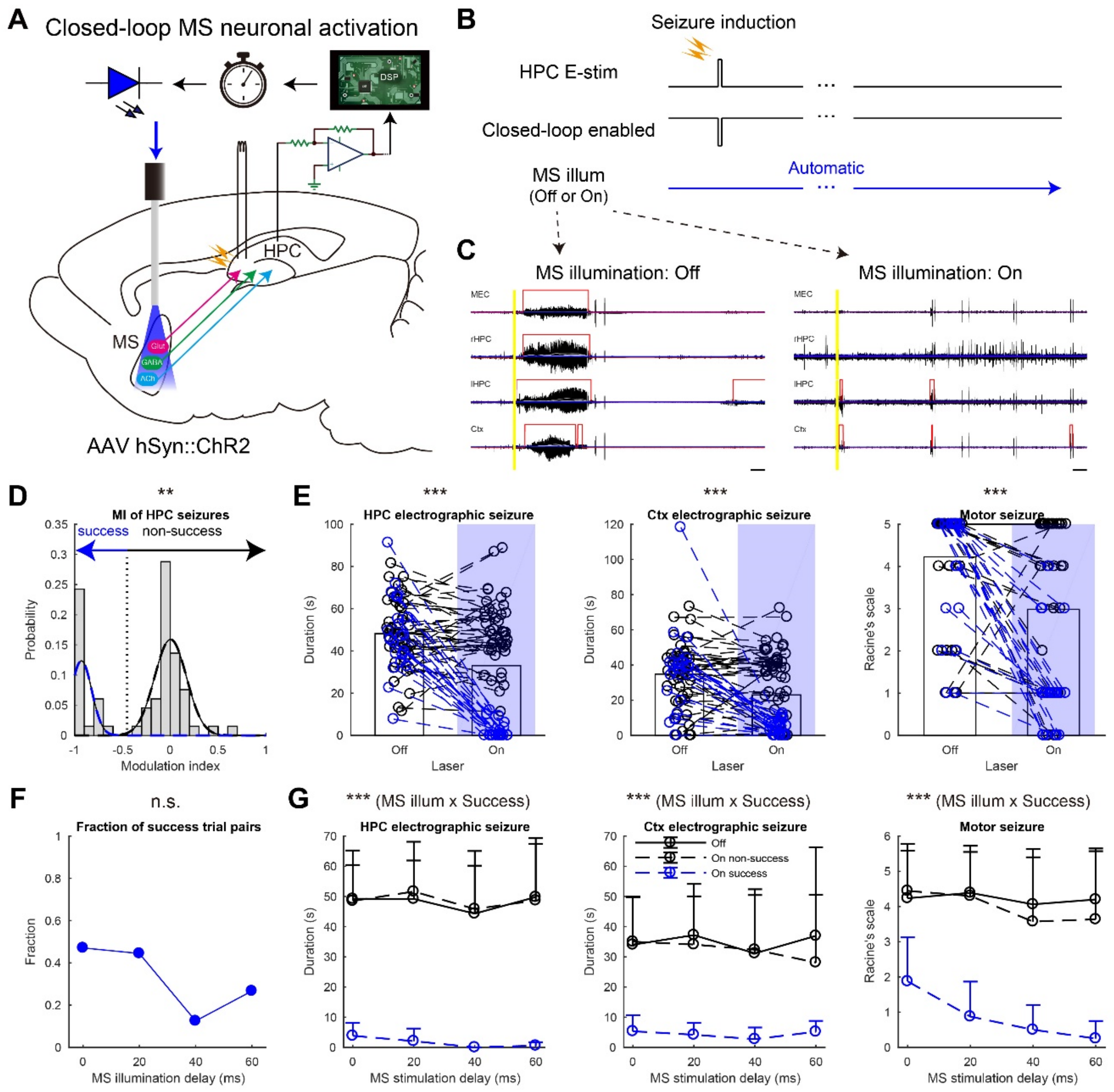


**Figure S3. Closed-loop seizure rhythm-driven optogenetic activation of MS neurons terminates seizures of HPC-origin, Related to Figure 3.**

(**A**) Schema of the experiment. (**B**) Closed-loop optogenetic seizure rhythm intervention. Seizures were induced by supra-threshold E-stim of the HPC commissure. Automatic detection of seizure waves was always turned on except during the seizure-induction time. Each detection triggers 30 ms-long MS illumination with a fixed delay. (**C**) Representative seizure waves w/wo closed-loop seizure rhythm MS illuminations. (**D**) Distribution of MIs. The distribution was fitted to a Gaussian mixture model and each trial pair was then clustered into success or non-success trial group. (**E**) Duration of HPC and Ctx electrographic seizures, and motor seizures with the clusters defined in (D): Blue, success trial pairs; Black, non-success trial pairs. Data with different delays were pooled. (**F**) Fractions of success trials as a function of illumination delays. (**G**) Illumination delay resolved representation of the data shown in (E). n = 132 trials from four rats. Other conventions for data presentation and statistical tests employed are the same as Figure 3. n.s., not significant; **$P < 0.01$; ***$P < 0.001$.

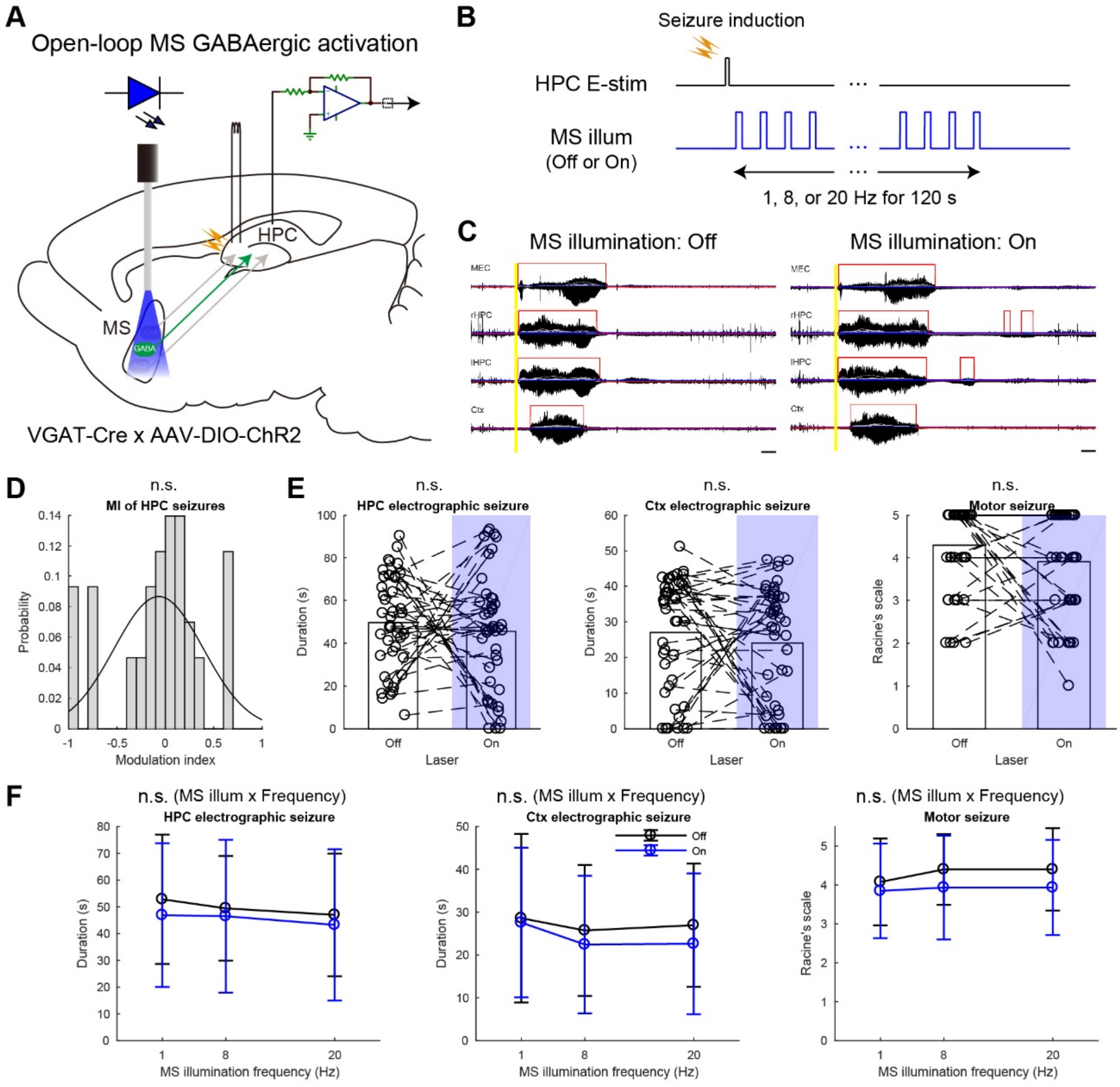


**Figure S4. Open-loop activation of MS GABAergic neurons does not alleviate seizures of HPC-origin, Related to Figure 4**

(**A**) Schema of the experiment. (**B**) Concept of open-loop responsive optogenetic seizure intervention. (**C**) Representative seizure waves w/wo open-loop responsive MS illuminations. (**D**) Distribution of MIs. (**E**) Duration of HPC and Ctx, and motor seizures. (**F**) Illumination frequency resolved representation of the data shown in (E). n = 86 trials from four rats. Other conventions for data presentation and statistical tests employed are the same as Figure S2. n.s., not significant.

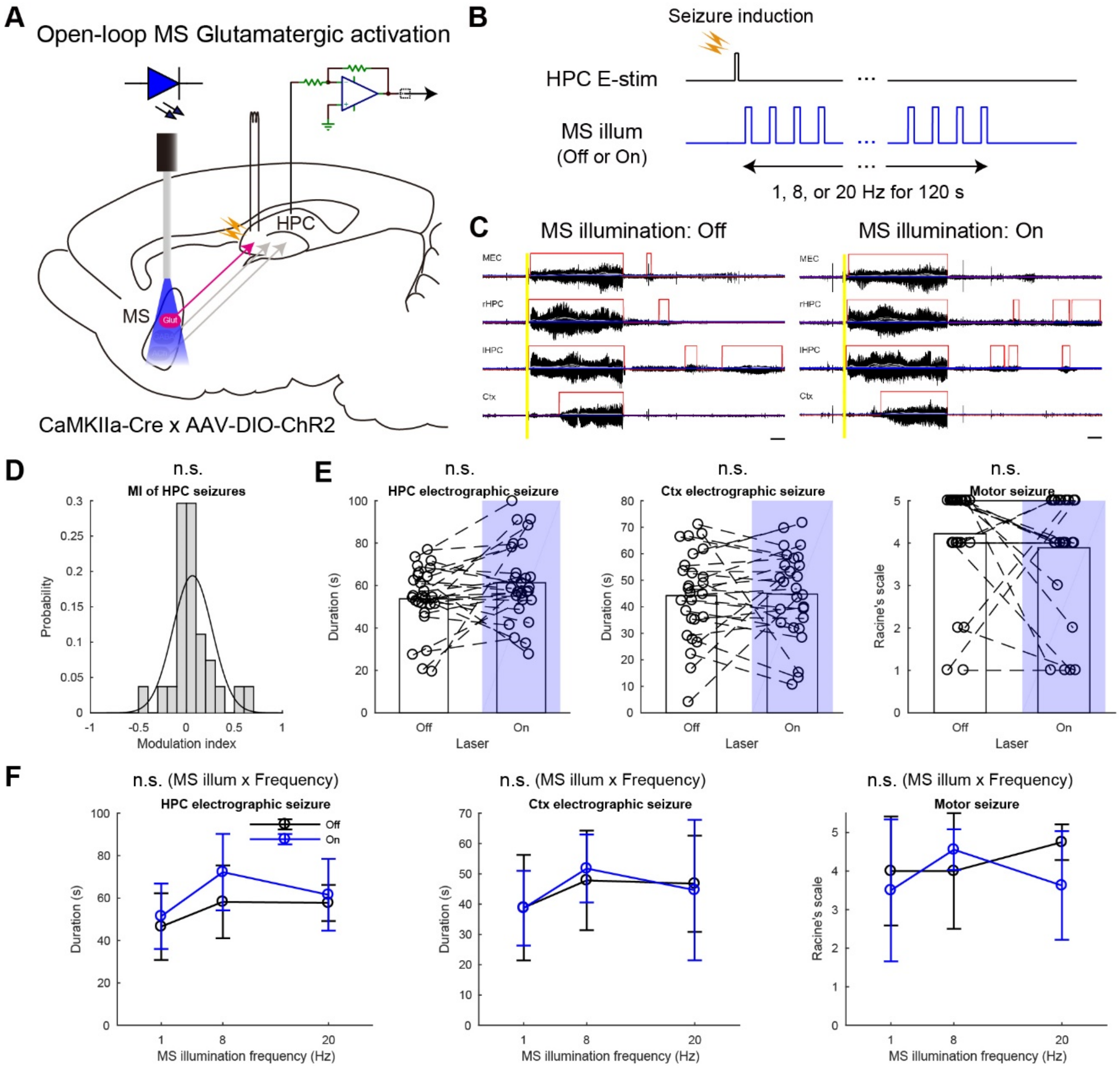


**Figure S5. Open-loop activation of MS Glut neurons does not alleviate seizures of HPC-origin, Related to Figure 5**

(**A**) Schema of the experiment. (**B**) Concept of open-loop responsive optogenetic seizure intervention. (**C**) Representative seizure waves w/wo open-loop responsive MS illuminations. (**D**) Distribution of MIs. (**E**) Duration of HPC and Ctx electrographic seizures, and motor seizures. (**F**) Illumination frequency resolved representation of the data shown in (E). n = 54 trials from two rats. Other conventions are the same as Figure S2. n.s., not significant.

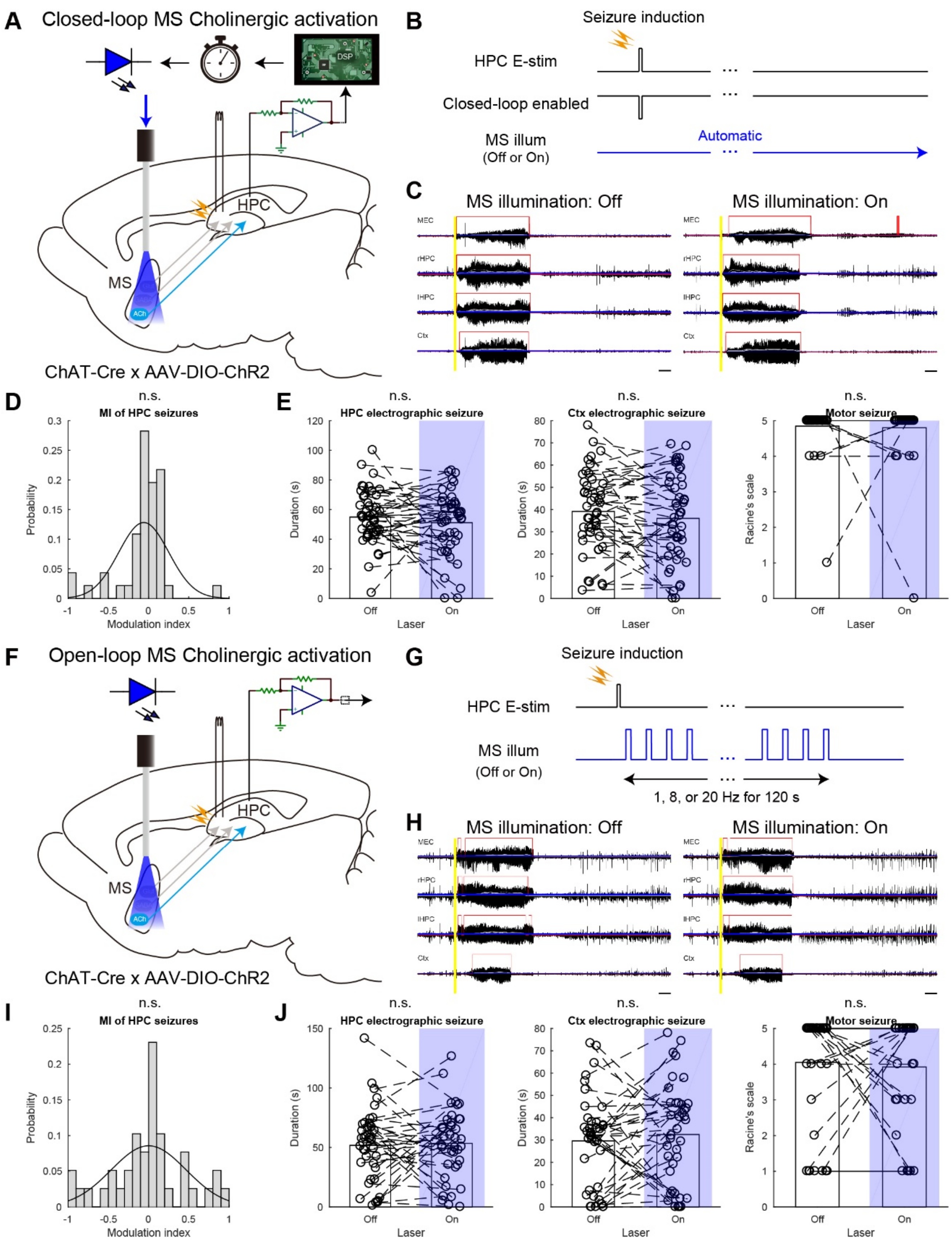


**Figure S6. Neither closed-loop nor open-loop activation of MS cholinergic neurons alleviates seizures of HPC-origin, Related to Figure 6.**

(**A**, **F**) Schemata of the experiments. (**B**) Concept of closed-loop seizure rhythm optogenetic intervention. (**C**, **H**) Representative seizure waves w/wo MS illuminations. (**D**, **I**) Distributions of MIs. Line plots represent fitted Gaussian models. (**E**, **J**) Duration of HPC and Ctx seizures, and motor seizures. Data of different illumination frequencies and delays were pooled. (**G**) Open-loop responsive optogenetic seizure intervention. n = 92 trials from four rats for (D, E) and 78 trials from three rats for (I, J). Other conventions for data presentation and statistical tests employed are the same as Figure S2. n.s., not significant.

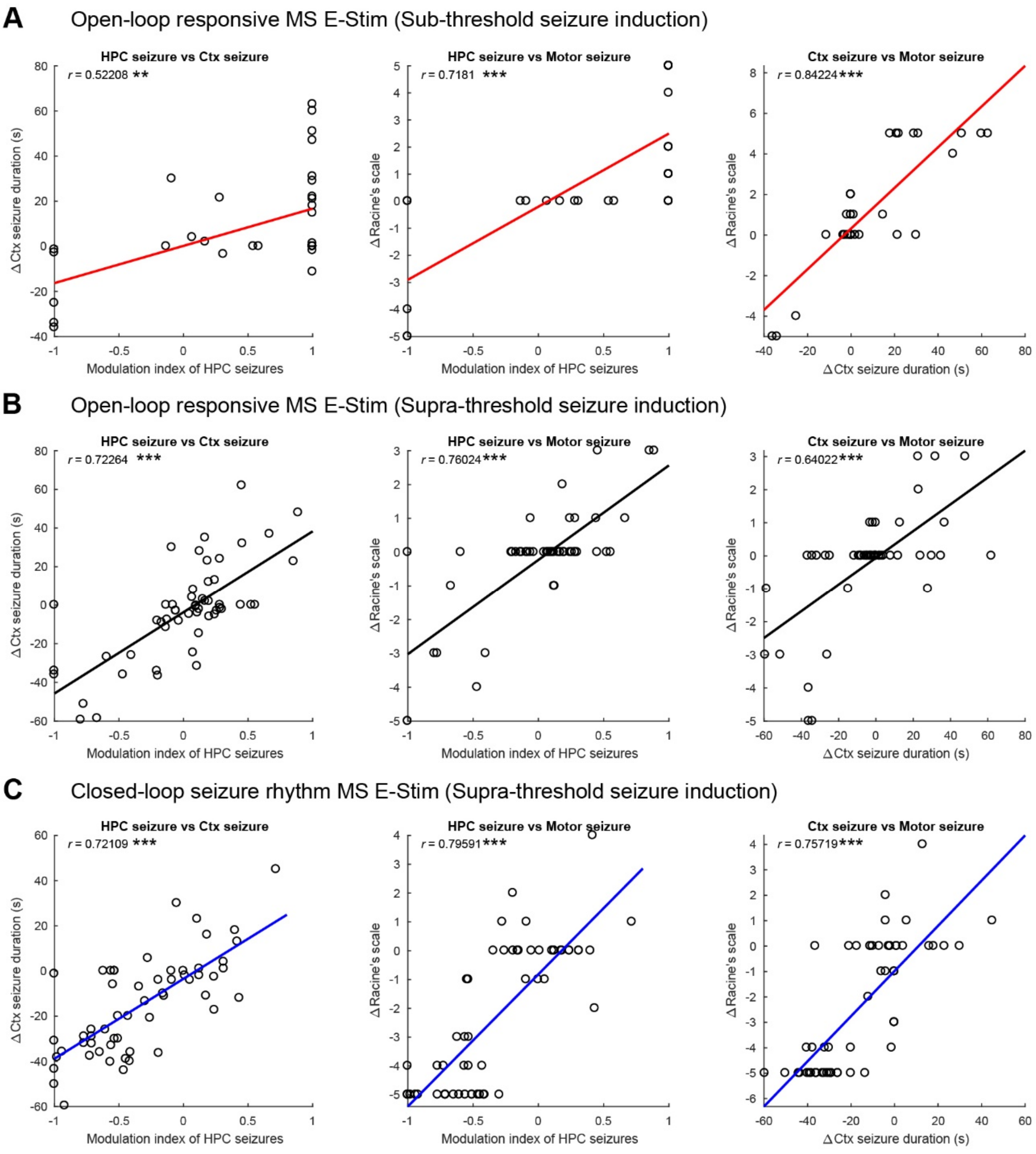

**Figure S7. Modulations of HPC seizures correlate with the changes of Ctx electrographic and motor seizures, related to Figure 7.**

(**A**) Correlations between MIs of duration of HPC seizures, changes of durations of Ctx seizures, and changes of severity of motor seizures w/wo open-loop responsive MS E-stim after sub-threshold seizure inductions. Each marker represents a seizure episode. Lines indicate regression lines. *r*: correlation coefficient. Data are the same as those plotted in Figure 2F, G. (**B**) The same conventions as in (A) but after supra-threshold seizure inductions. Data are the same as those plotted in Figure 2D, E. (**C**) The same convention as in (A) but w/wo closed-loop seizure rhythm-driven MS E-stim. Data are the same as those plotted in Figure 3E, F. Statistical significance was tested by Spearman's rank correlation test. **$P < 0.01$; ***$P < 0.001$.

**Table S1. Statistical table**

Provided as a separate MS Excel file.

**Table S2. Key Resource Table**

| REAGENT or RESOURCE | SOURCE | IDENTIFIER |
|---|---|---|
| Antibodies | | |
| Mouse monoclonal anti-NeuN | Millipore | Cat# MAB377; RRID:AB_2298772 |
| Rabbit polyclonal anti-GAD67/65 | Frontier Institute | Cat# GAD-Rb; RRID:AB_2571698 |
| Mouse monoclonal anti-glutamate | ImmunoStar | Cat# 22523; RRID:AB_572244 |
| Goat polyclonal anti-ChAT | Millipore | Cat# AB144P; RRID:AB_2079751 |
| Rabbit polyclonal anti-cFos | Millipore | Cat# ABE457; RRID:AB_2631318 |

| | | |
|---|---|---|
| Mouse monoclonal anti-PV | Swant | Cat# 235; RRID: AB_10000343 |
| Alexa Fluor 488 goat anti-mouse | Thermo Fisher Scientific | Cat# A-11029, RRID:AB_2534088 |
| Alexa Fluor 633 goat anti-rabbit | Thermo Fisher Scientific | Cat# A-21071, RRID:AB_2535732 |
| Alexa Fluor 647 donkey anti-mouse | Thermo Fisher Scientific | Cat# A-31571; RRID:AB_162542 |
| Alexa Fluor 633 donkey anti-goat | Thermo Fisher Scientific | Cat# A-21082; RRID:AB_2535739 |
| Alexa Fluor 488 donkey anti-goat | Thermo Fisher Scientific | Cat# A-11055; RRID:AB_2534102 |
| Alexa Fluor 555 donkey anti-rabbit | Thermo Fisher Scientific | Cat# A-31572, RRID:AB_162543 |
| Bacterial and Virus Strains | | |
| AAV5-hSyn-hChR2(H134R)-mCherry | UNC Vector Core | N/A |
| AAV5-EF1α-DIO-hChR2(H134R)-mCherry | UNC Vector Core | N/A |
| Chemicals, Peptides, and Recombinant Proteins | | |
| Colchicine | Sigma-Aldrich | Cat# C9754 |
| Urethane | Sigma-Aldrich | Cat# U2500 |
| Paraformaldehyde | Sigma-Aldrich | Cat# P6148 |
| Glutaraldehyde | Sigma-Aldrich | Cat# 354400 |
| DAPI | Sigma-Aldrich | Cat# D8417 |
| NeuroTrace 435/455 blue-fluorescent Nissl stain | Thermo Fisher Scientific | Cat# N-21479 |
| DiD | Thermo Fisher Scientific | Cat# D307 |
| Experimental Models: Organisms/Strains | | |
| Rat: LE-CamKIIa-Cre$^{tm1sage}$ | Horizon Discovery | RRID:RGD_12905032 |
| Rat: LE-Slc32a1$^{tm1(cre)Sage}$ | Horizon Discovery | RRID:RGD_12905033 |

| | | |
|---|---|---|
| Rat: LE-Tg(ChAT-Cre)5.1Deis | Witten et al., 2011 (1) | RRID:RGD_10401204 |
| Oligonucleotides | | |
| Primer: Cre Forward: 5'-ACCTGATGGACATGTTCAGGGATCG-3' | Iwasato et al, 2004 (2) | N/A |
| Primer: Cre Reverse: 5'-TCCGGTTATTCAACTTGCACCATGC-3' | Iwasato et al, 2004 (2) | N/A |
| Software and Algorithms | | |
| MATLAB R2017a | Mathworks | RRID:SCR_001622 |
| Neuroscope | Hazan et al., 2006 (3) | RRID:SCR_002455 |
| ZEN Digital Imaging for Light Microscopy | Carl Zeiss | RRID:SCR_013672 |
| ImageJ 1.49v | NIH | RRID:SCR_003070 |
| Klusta | Kadir et al., 2014 (4) | https://github.com/kwikteam/klusta |
| KlustaViewa | Rossant et al., 2016 (5) | https://github.com/klusta-team/klustaviewa/ |
| Circular Statistics Toolbox | Berens et al., 2009 (6) | RRID:SCR_016651 |
| Other | | |
| HML Insulated Tungsten 99.95% wire | California Fine Wire | Cat# CFW2044436 |

### References

1. Witten IB, Steinberg EE, Lee SY, Davidson TJ, Zalocusky KA, Brodsky M, et al. Recombinase-driver rat lines: tools, techniques, and optogenetic application to dopamine-mediated reinforcement. Neuron. 2011;72(5):721-33.

2. Iwasato T, Nomura R, Ando R, Ikeda T, Tanaka M, Itohara S. Dorsal telencephalon-specific expression of Cre recombinase in PAC transgenic mice. Genesis. 2004;38(3):130-8.

3. Hazan L, Zugaro M, Buzsáki G. Klusters, NeuroScope, NDManager: A free software suite for neurophysiological data processing and visualization. J Neurosci Methods. 2006;155(2):207-16.

4. Kadir SN, Goodman DF, Harris KD. High-Dimensional Cluster Analysis with the Masked EM Algorithm. Neural comput. 2014;26(11):2379-94.

5. Rossant C, Kadir SN, Goodman DF, Schulman J, Hunter ML, Saleem AB, et al. Spike sorting for large, dense electrode arrays. Nat Neurosci. 2016;19(4):634-41.

6. Berens P. CircStat: A MATLABToolbox for Circular Statistics. J Statisticcal Software. 2009;31(10):10.
