## Supplementary material for "Closed-loop stimulation of the medial septum terminates epileptic seizures": Dataset figures 1-9

### Dataset figure 1

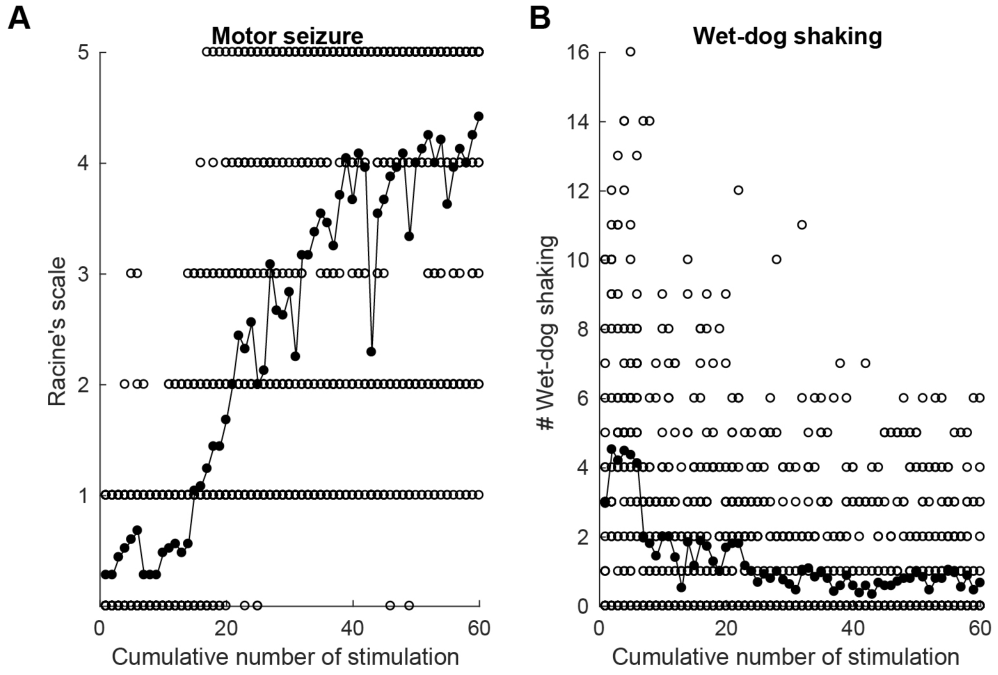


**Development of motor seizures over 10-day-long kindling processes.** Severity of each motor seizure induced by electrical stimulation of the hippocampal commissure was evaluated using the Racine's scale (A) and the number of wet-dog shaking behaviors (B). The Racine's scale: 1, Mouth and facial movements; 2, Head nodding; 3, Forelimb clonus; 4, Rearing; 5, Rearing and falling. Six stimulation were conducted per day. Each stimulation consists of 62.5 Hz bipolar 120 constant current pulses on the hippocampal commissure at the intensity by which after discharges in the hippocampal local field potentials are stably observed: ±20–200 μA typically. Each open circle represents the Racine's scale or number of wet-dog shaking behaviors of each trial. Each closed circle represents the mean value over rats. Number of rats: 25.

### Dataset figure 2

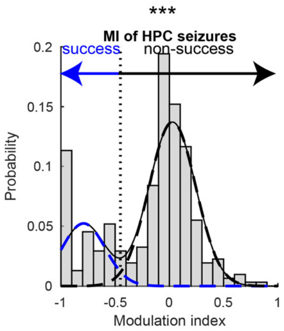


**Determination of the global threshold of modulation index for success and non-success trials.** Distribution of modulation index (MI) with the MS illumination. The distribution was fitted with a Gaussian mixture model to cluster each trial pair into success or non-success trial group. The global threshold (dotted vertical line) was determined as a MI value by which the two Gaussian model curves cross. Data of the whole closed-loop interventions were pooled from Figures 3E, 4E, 5F, and S3D. The data represent 618 trials from 18 rats in total. Statistical significance was tested by two-sample Kolmogorov-Smirnov for asymmetry (skewness) of MI distribution. ***$P < 0.001$.

### Dataset figure 3

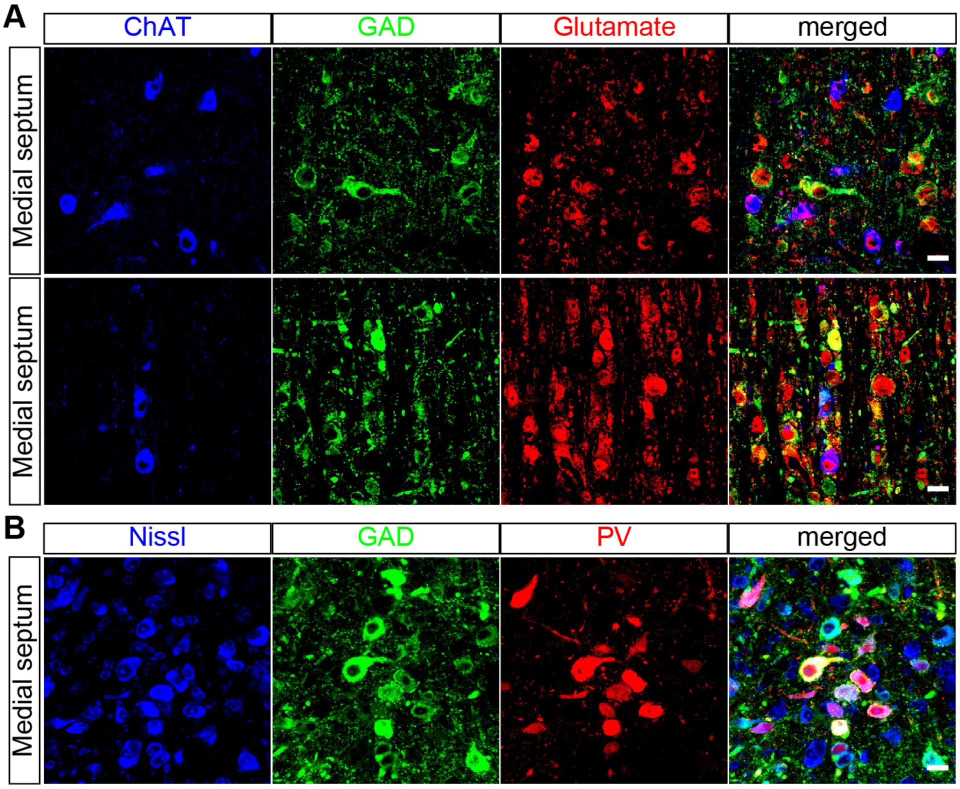


**Neuronal populations in the medial septum revealed by immunohistochemistry**. (**A**) Cholinergic, GABAergic, and glutamatergic neurons supposed to be ChAT-positive, GAD-positive, and ChAT-GAD-double negative populations, respectively. (**B**) PV-positive neuronal population is a sub-subpopulation of GAD-positive neuronal populations. Scale bars: 20 µm. ChAT, choline acetyltransferase; GAD, glutamate decarboxylase; PV, parvalbumin.

### Dataset figure 4

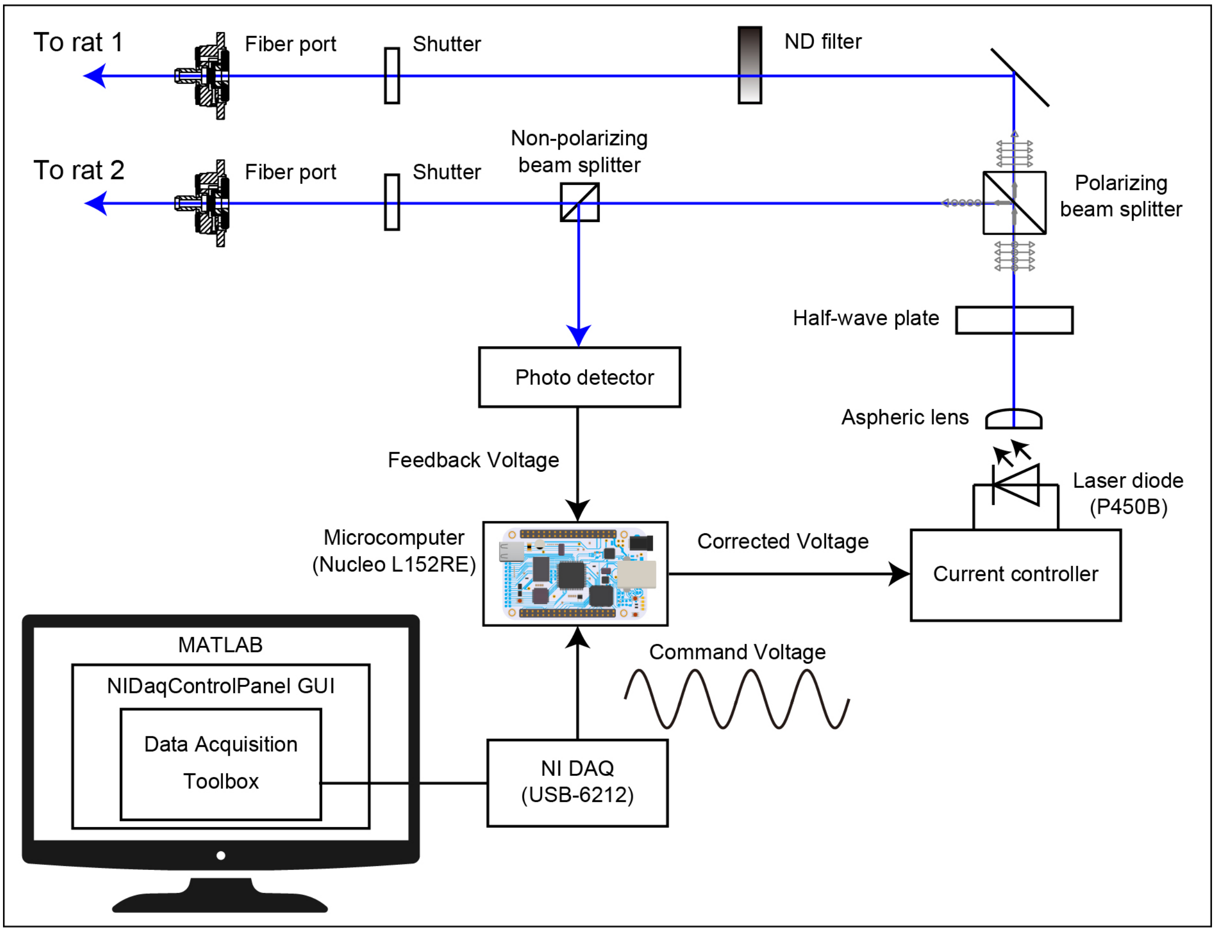


**Light source for optogenetic stimulation.** Laser diode with 450 nm wavelength was driven by a current controller with an external analog modulation. External command voltages were provided from a NI DAQ board or multichannel stimulator. The power of the 450 nm laser ray (up to 130 mW) was collimated and directed to port 1 and/or port 2 depending on its polarity using a half-wave plate and polarizing beam splitter. Combined with manual shutters, optogenetic stimulation was conducted on port 1 or 2 alternatingly to increase throughput of the experiments. Optical output power of the laser diode was stabilized with a self-made feedback system driven by a photo detector and a microcomputer (https://github.com/yuichi-takeuchi/LaserDiodeStabilizer).

### Dataset figure 5

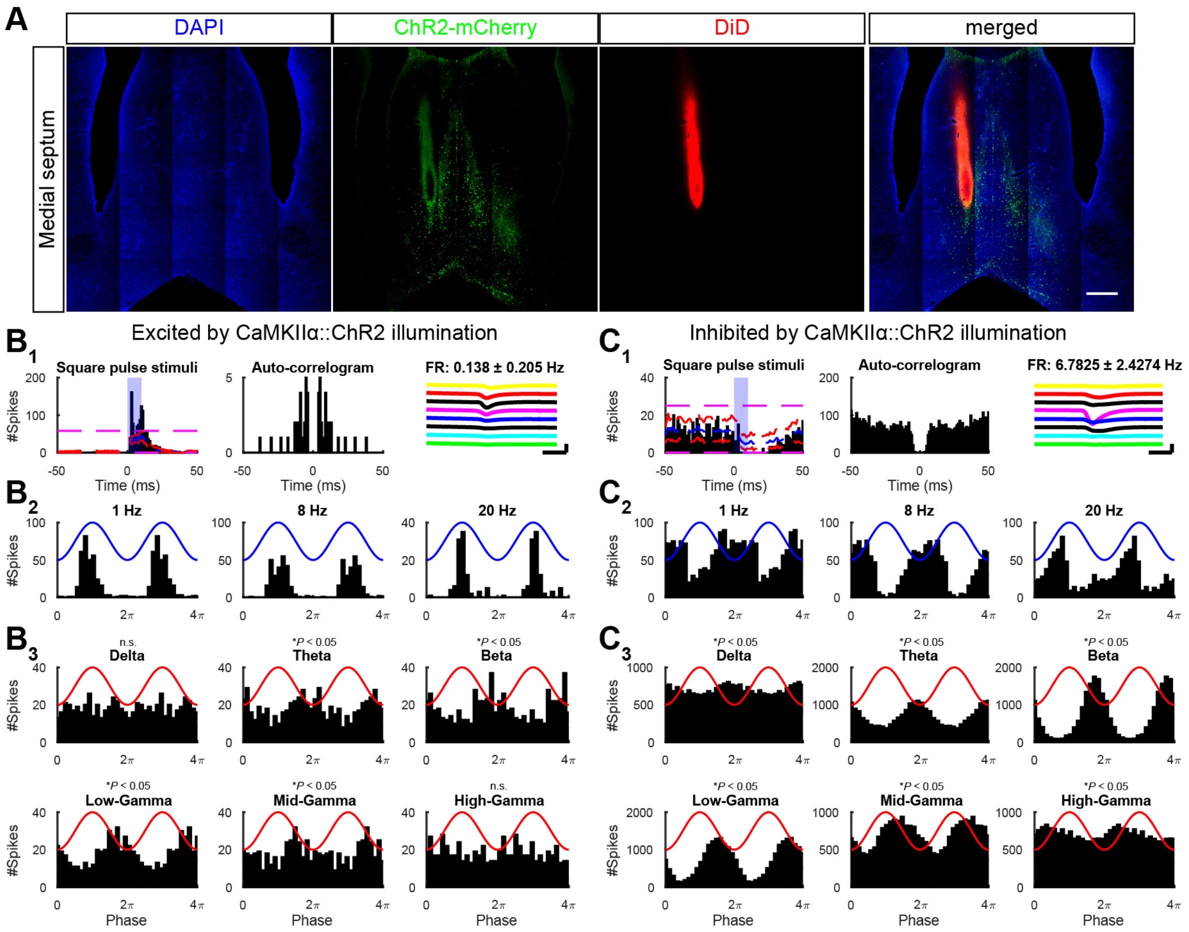


**Examples of optogenetic manipulation of neuronal activity in the medial septum.** (**A**) A track of silicon probe recording electrode in the medial septum. This recording was made from the medial septum (MS) of an urethane-anesthetized CaMKIIα::ChR2 rat. Scale bar: 0.5 mm. (**B**, **C**) Examples of excited and inhibited septal neurons by blue-light illumination in the MS of a CaMKIIα::ChR2 rat. ($B_1$, $C_1$) Peri-stimulus time histogram (PSTH), auto-correlogram, and waveforms. Dashed pink, red, and blue lines in PSTHs represent global significant lines, local significant lines, and mean counts after shuffling. Blue patches represent optical illumination. Waveform scalebars: 0.2 mV, 0.5 ms. ($B_2$, $C_2$) Excitatory and inhibitory modulated firing in response to sinusoidal blue light illumination at 1, 8, and 20 Hz. ($B_3$, $C_3$) Phase-modulation of unit firing by different frequency bands of local field potentials in the MS. Statistical significance for phase-modulation of unit was tested by Rayleigh's test. n.s., not significant; $*P < 0.05$. DiD, 1,1'-dioctadecyl-3,3,3',3'-tetramethylindodicarbocyanine perchlorate.

### Dataset figure 6

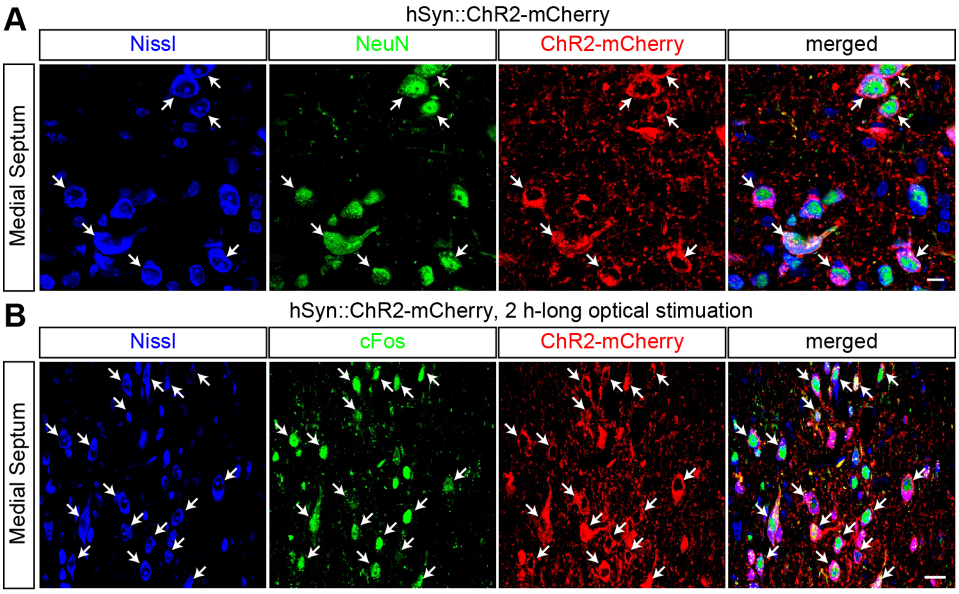


**Expression and functional test of ChR2 in MS of hSyn::ChR2 rats**. (**A**) Channelrhodopsin-2 (ChR2) was transduced into the MS of wild-type Long-Evans rats with an AAV5-hSyn-ChR2-mCherry. ChR2-mCherry signals were colocalized NeuN positive neurons. Each arrow indicates the colocalization. Scale bar: 10 µm. (**B**) Expression of cFos, an immediate early gene, after two-hour-long 20 Hz sine wave illumination of the AAV5-hSyn-ChR2-mCherry-injected MS. Immunohistochemically visualized cFos was colocalized with ChR2 expressing neurons. Each arrow indicates the colocalization. Scale bar: 20 µm.

### Dataset figure 7

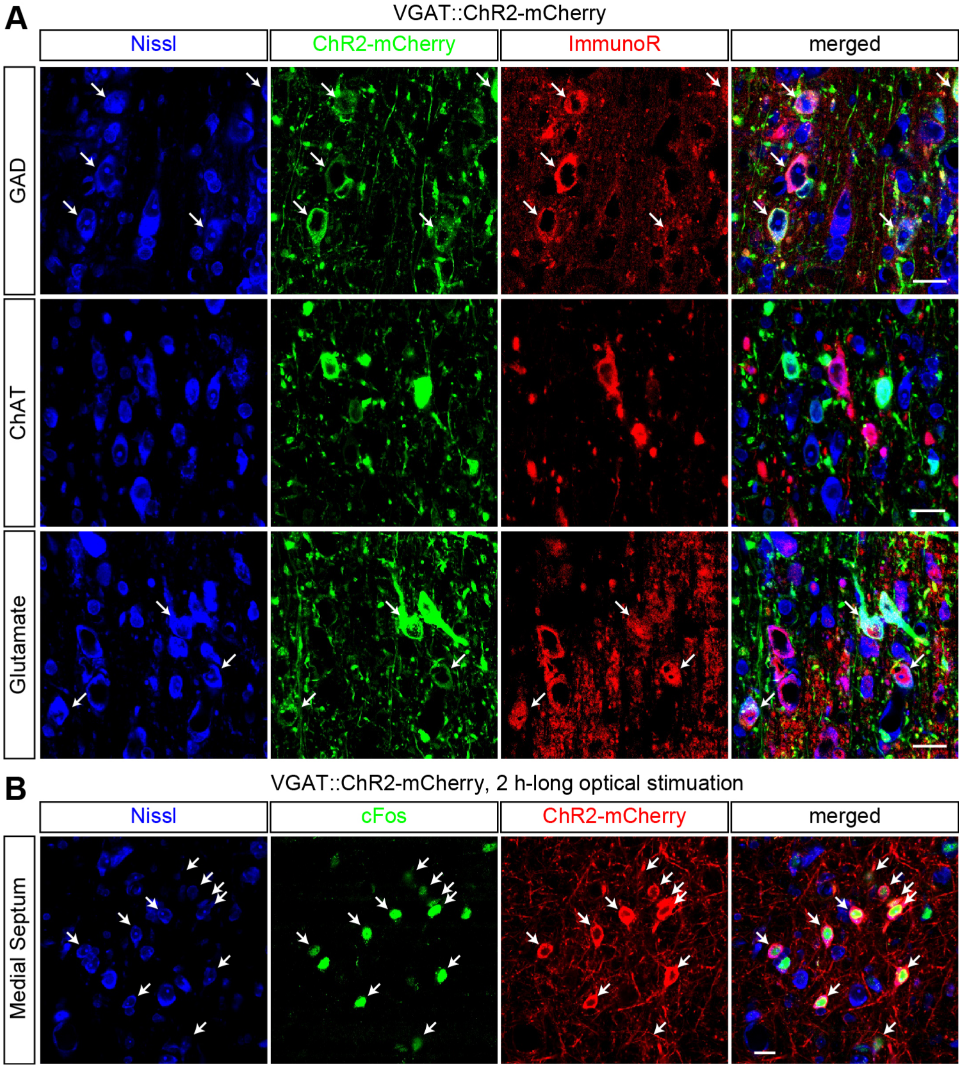


**Expression and functional test of ChR2 in MS of VGAT::ChR2 rats.** (**A**) ChR2 was transduced into the MS of VGAT-Cre rats with an AAV5-EF1α-DIO-ChR2-mCherry. ChR2 expression was colocalized with immunohistochemically visualized GAD and glutamate, but not with ChAT. Each arrow indicates the colocalization of ChR2 and GAD- or glutamate-positive neurons. Scale bars: 20 µm. (**B**) Expression of cFos, an immediate early gene, after two-hour-long 20 Hz sine wave illumination of the AAV5-DIO-ChR2-injected MS of VGAT::ChR2 rats. Immunohistochemically visualized cFos was colocalized with ChR2 neurons. Each arrow indicates the colocalization. Scale bar: 20 µm. ChAT, choline acetyltransferase; GAD, glutamate decarboxylase; ImmunoR, immunohistochemical reaction.

### Dataset figure 8

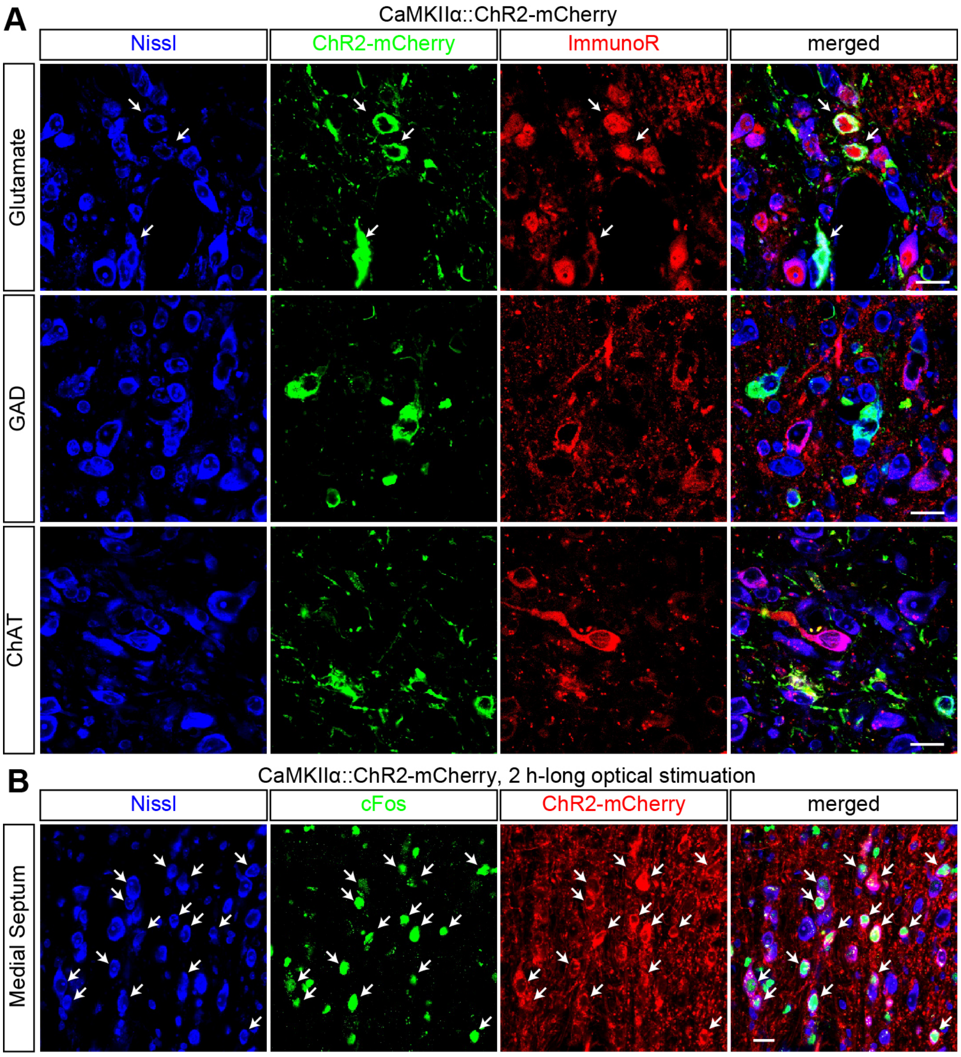


**Expression and functional test of ChR2 in MS of CaMKIIα::ChR2 rats.** (**A**) ChR2 was transduced into the MS of CaMKIIα-Cre rats with an AAV5-EF1α-DIO-ChR2-mCherry. ChR2 expression was colocalized with immunohistochemically visualized of glutamate, but not with GAD or ChAT. Each arrow indicates the colocalization. Scale bars: 20 µm. (**B**) Expression of cFos, an immediate early gene, after two-hour-long 20 Hz sine wave illumination of the AAV5-DIO-ChR2-injected MS of CaMKIIα::ChR2 rats. Immunohistochemically visualized cFos was colocalized with ChR2 expressing neurons. Each arrow indicates the colocalization. Scale bar: 20 µm. ChAT, choline acetyltransferase; GAD, glutamate decarboxylase; ImmunoR, immunohistochemical reaction.

### Dataset figure 9

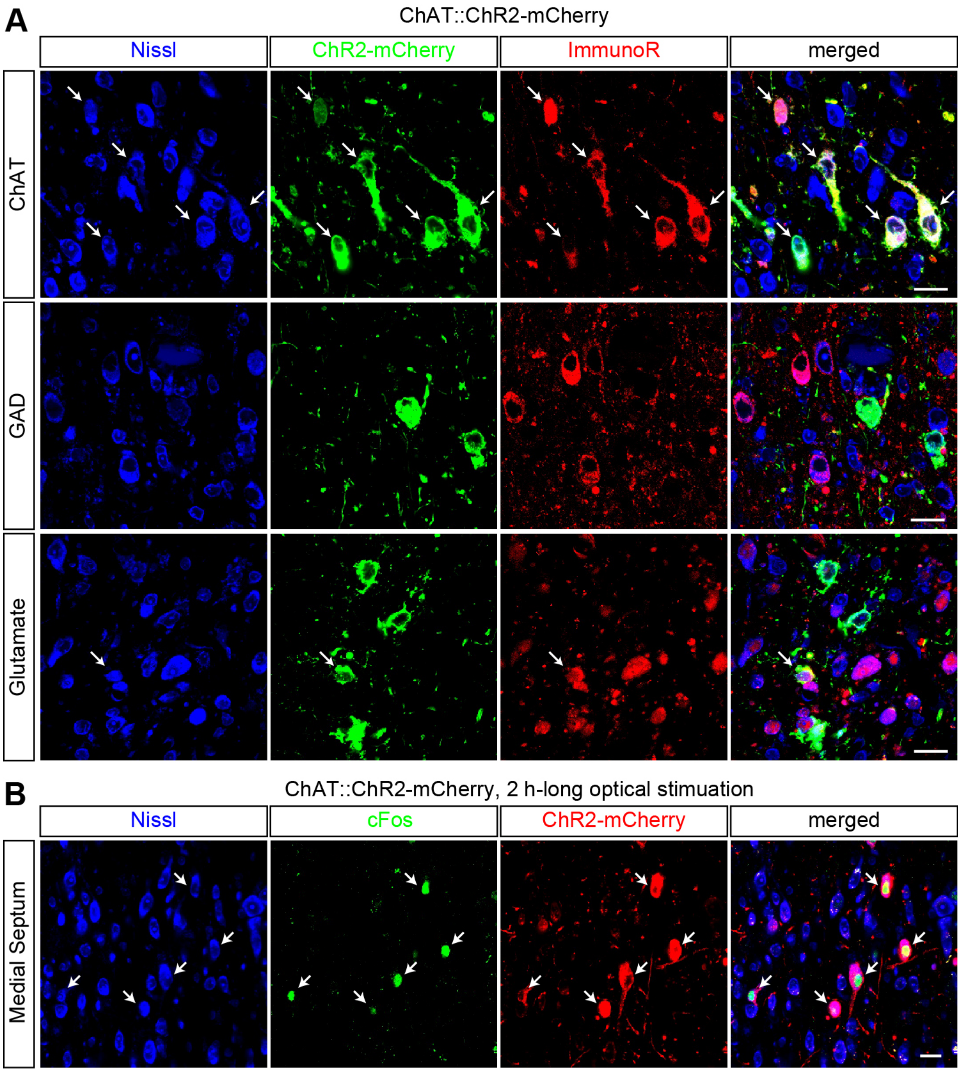


**Expression and functional test of ChR2 in MS of ChAT::ChR2 rats.** (**A**) ChR2 was transduced into the MS of ChAT-Cre rats with an AAV5-EF1α-DIO-ChR2-mCherry. ChR2 expression was colocalized with immunohistochemically visualized ChAT and glutamate, but not with GAD. Each arrow indicates the colocalization of ChR2 and ChAT- or glutamate-positive neurons. Scale bars: 20 µm. (**B**) cFos immediate early gene expression after two-hour-long 20 Hz sine wave illumination of the AAV5-DIO-ChR2-injected MS of ChAT::ChR2 rats. Immunohistochemically visualized cFos was colocalized with ChR2 expressing neurons. Each arrow indicates the colocalization. Scale bar: 20 µm. ChAT, choline acetyltransferase; GAD, glutamate decarboxylase; ImmunoR, immunohistochemical reaction.
